# 2-Thiouridine is a broad-spectrum antiviral nucleoside analogue against positive-strand RNA viruses

**DOI:** 10.1101/2022.12.14.520006

**Authors:** Kentaro Uemura, Haruaki Nobori, Akihiko Sato, Shinsuke Toba, Shinji Kusakabe, Michihito Sasaki, Koshiro Tabata, Keita Matsuno, Naoyoshi Maeda, Shiori Ito, Mayu Tanaka, Yuki Anraku, Shunsuke Kita, Mayumi Ishii, Kayoko Kanamitsu, Yasuko Orba, Yoshiharu Matsuura, William W. Hall, Hirofumi Sawa, Hiroshi Kida, Akira Matsuda, Katsumi Maenaka

**Author notes:** Corresponding authors: Akihiko Sato;, Akira Matsuda;, Katsumi Maenaka (lead contact).

## Abstract

Severe acute respiratory syndrome coronavirus 2 (SARS-CoV-2) infection causes significant morbidity and mortality worldwide, seriously impacting not only human health but also the global economy. Furthermore, over 1 million cases of newly emerging or re-emerging viral infections, specifically dengue virus (DENV), are known to occur annually. Because no virus-specific and fully effective treatments against these and many other viruses have been approved, they continue to be responsible for large-scale epidemics and global pandemics. Thus, there is an urgent need for novel, effective therapeutic agents. Here, we identified 2-thiouridine (s2U) as a broad-spectrum antiviral nucleoside analogue that exhibited antiviral activity against SARS-CoV-2 and its variants of concern, including the Delta and Omicron variants, as well as a number of other positive-sense single-stranded RNA (ssRNA+) viruses, including DENV. s2U inhibits RNA synthesis catalyzed by viral RNA-dependent RNA polymerase, thereby reducing viral RNA replication, which improved the survival rate of mice infected with SARS-CoV-2 or DENV in our animal models. Our findings demonstrate that s2U is a potential broad-spectrum antiviral agent not only against SARS-CoV-2 and DENV but other ssRNA+ viruses.

## Main text

Severe acute respiratory syndrome coronavirus 2 (SARS-CoV-2; family: *Coronaviridae*) is responsible for the ongoing pandemic of coronavirus disease 2019 (COVID-19), and the number of cases since December 2019 has been estimated at over 648 million^1^. Likewise, many newly emerging or re-emerging viral infections have occurred in the 21st century and also continue to seriously impact human health. For instance, dengue virus (DENV; family: *Flaviviridae*) causes large-scale epidemics in Asia, with 390 million cases annually^2^, and Chikungunya virus (CHIKV; family: *Togaviridae*), which is endemic in the equatorial regions, causes at least 3 million infections annually^3^. The genomes of these viruses are composed of positive-sense single-stranded RNA (ssRNA+). Although extensive drug discovery research against these viruses has been undertaken, no virus-specific and fully effective treatments have been approved. With respect to SARS-CoV-2, emergency use authorization (EUA) of neutralizing antibodies has significantly contributed to suppressing disease severity, these are expensive and only effective for the fusion step of SARS-CoV-2 infection. Therefore, the development of novel effective therapeutic agents, particularly small-molecule compounds, against COVID-19 and many other viral diseases is essential.

Because licensed antiviral therapeutic agents are limited, broad-spectrum antiviral agents are critical for the control of emerging viral diseases. Ribavirin, which was developed in the 1970s, is a broad-spectrum antiviral agent effective against both RNA and DNA viruses^4–6^, and it has been reported to be clinically beneficial in the treatment of several viral diseases^7–9^. In the current COVID-19 situation, remdesivir^10^ and molnupiravir^11^ were developed as anti-SARS-CoV-2 agents and have received EUA. Remdesivir is an adenosine analogue that exhibits antiviral activities against the *Filoviridae, Paramyxoviridae, Pneumoviridae, Flaviviridae* and *Coronaviridae* families^12,13^. Molnupiravir is a cytidine analogue that exhibits antiviral activities against influenza virus (IFV), respiratory syncytial virus (RSV) and SARS-CoV-2^14–16^. The molecular target of these nucleoside analogues or their prodrugs is viral RNA-dependent RNA polymerase (RdRp), and they inhibit viral RNA replication by inhibiting RdRp activity or by introducing a mutation into the viral genome during replication^17–19^. RdRp is indispensable for the replication and transcription of viral genomes, and the core structural features of viral RdRps are functionally essential and conserved across a wide range of viruses^20–22^. Thus, viral RdRp is a promising target for the development of broad-spectrum inhibitors of viruses with this.

Here, we report the discovery of a broad-spectrum antiviral nucleoside analogue, 2-thiouridine (s2U), using phenotypic screening. Furthermore, we have successfully demonstrated the antiviral activity of s2U against several ssRNA+ viruses, including SARS-CoV-2 and DENV serotype 2 (DENV2), *in vitro* and *in vivo.*

### s2U is a broad-spectrum inhibitor of ssRNA+ viruses

We initially performed cell-based anti-DENV2 screening of 753 nucleoside analogues in a compound library from Hokkaido University. We identified a number of hit compounds and further confirmed their antiviral effects against all serotypes of DENV and flaviviruses, including Zika virus (ZIKV), yellow fever virus (YFV), Japanese encephalitis virus (JEV) and West Nile virus (WNV) (Extended Data Fig. 1a). We have clearly identified s2U (Fig. 1a) as a nucleoside analogue with strong antiviral activity against all eight tested flaviviruses (Table 1). The antiviral activity of s2U was markedly higher than that of ribavirin and favipiravir, which are known to have anti-flavivirus activity^23,24^ (Table 1, Extended Data Table 1).

**Fig. 1.**
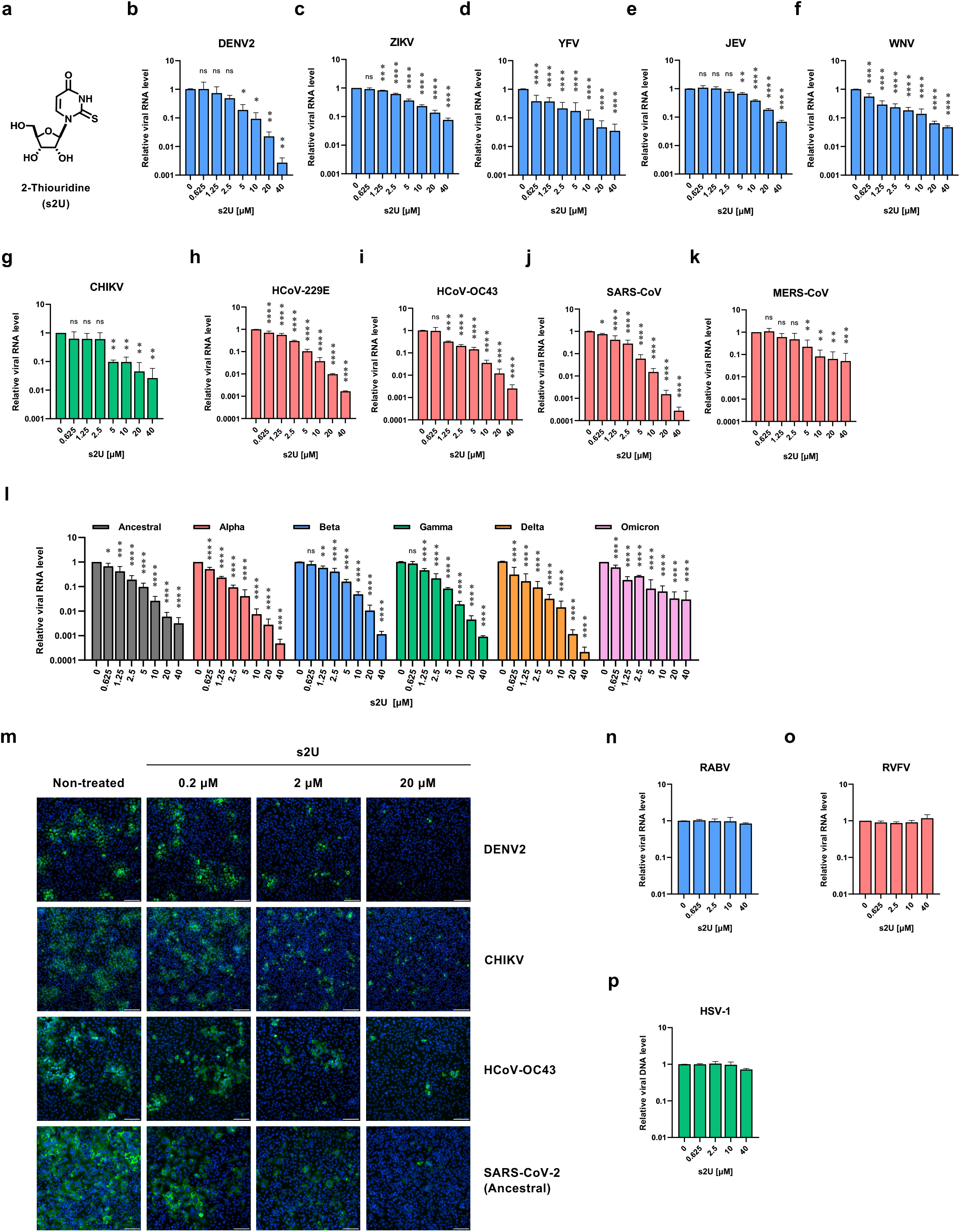
Broad-spectrum antiviral activities of s2U against several RNA viruses. **a**, Chemical structure of 2-thiouridine (s2U). **b–l**, Dose-response inhibition of DENV2 (**b**), ZIKV (**c**), YFV (**d**), JEV (**e**), WNV (**f**), CHIKV (**g**), HCoV-229E (**h**), HCoV-OC43 (**i**), SARS-CoV (**j**), MERS-CoV (**k**) and several SARS-CoV-2 variants (**l**) by s2U. Cell lysates were collected for viral RNA determination, and viral RNA levels were determined relative to *ACTB* or *18S rRNA* transcripts. **m**, Dose-response inhibition of viral protein expression in the DENV2-, CHIKV-, HCoV-OC43- and SARS-CoV-2-infected cells. Cells were stained with viral-specific antibodies (green) and counterstained with Hoechst 33342 nuclear dye (blue). Scale bars indicate 200 μm. **n–o**, s2U did not inhibit RABV (**n**) and RVFV (**o**) virus replication. Cell lysates were collected for viral RNA determination; viral RNA levels were determined relative to *ACTB* or *18S rRNA* transcripts. **p**, s2U did not inhibit HSV-1 virus replication. Cell lysates were collected for viral DNA determination; viral DNA levels were determined relative to *ACTB* transcripts. Data are presented as mean values of biological triplicates from one of the experiments, and error bars indicate standard deviation (SD). Statistically significant differences were determined using a one-way ANOVA followed by Dunnett’s multiple comparisons test to compare with non-treated cells; * *p* < 0.01, ** *p* < 0.005, *** *p* < 0.0005 and **** *p* < 0.0001.

**Table 1.** Antiviral activity of s2U against various RNA and DNA viruses. Antiviral assays were carried out as described in Supplementary Table 1. EC_50_: 50% effective concentration. EC_90_: 90% effective concentration. CC_50_: 50% cytotoxic concentration. a: Selectivity index (SI) is calculated from the ratio of CC_50_/EC_50_. b: EC_50_ values represent mean values ± SEM from at least three independently performed experiments (n = 2). c: EC_50_ values represent mean values from at least three independently performed experiments (n = 2). d: EC_50_ and EC_90_ values represent mean values from a single experiment with biological triplicates (see Extended Data Fig. 2). e: Fold change is calculated from the ratio of rgNS5-G605V/rgWT.

| Screening Results |  |  |  |  |  |  |  |  |  |
| --- | --- | --- | --- | --- | --- | --- | --- | --- | --- |
| Virus | Strain | Cell | EC<sub>50</sub> (μM) | CC<sub>50</sub> (μM) | SI <sup>a</sup> |  |  |  |  |
| DENV1 | D1/hu/PHL/10-07 | BHK-21 | 2.6 | >400 | >154 |  |  |  |  |
| DENV2 | D2/hu/INDIA/09-74 | BHK-21 | 0.58 | >400 | >690 |  |  |  |  |
| DENV3 | D3/hu/Thailand/00-40 | BHK-21 | 2.1 | >400 | >190 |  |  |  |  |
| DENV4 | D4/hu/Solomon/09–11 | BHK-21 | 1.8 | >400 | >222 |  |  |  |  |
| ZIKV | MR766 | BHK-21 | 5.0 | >400 | >80 |  |  |  |  |
| YFV | 17D-204 | BHK-21 | 2.9 | >400 | >138 |  |  |  |  |
| JEV | Beijing-1 | BHK-21 | 4.8 | >400 | >83 |  |  |  |  |
| WNV | NY99 | BHK-21 | 5.3 | >400 | >75 |  |  |  |  |
| MTT / Resazurin Assay Results |  |  |  |  |  |  |  |  |  |
| Virus | Strain | Cell | EC<sub>50</sub> (μM) <sup>b</sup> |  | Virus | Strain | Cell | EC<sub>50</sub> (μM) <sup>c</sup> |  |
| DENV2 | D2/hu/INDIA/09-74 | BHK-21 | 1.5 ± 0.10 |  | RABV | HEP | BHK-21 | >100 |  |
| ZIKV | MR766 | BHK-21 | 3.8 ± 0.96 |  | LACV | ATCC VR-1834 | MDBK | >100 |  |
| YFV | 17D-204 | BHK-21 | 3.2 ± 1.4 |  | LPHV | 11SB17 | KB | >100 |  |
| JEV | Beijing-1 | VeroE6 | 7.6 ± 1.3 |  | LCMV | Armstrong | KB | >100 |  |
| WNV | NY99 | BHK-21 | 8.6 ± 1.8 |  | JUNV | Candid #1 | 293T | >100 |  |
|  |  | VeroE6 | 4.0 ± 0.24 |  | SFTSV | ArtLN/2017 | MDCK | >100 |  |
| CHIKV | SL10571 | BHK-21 | 3.8 ± 0.12 |  | RVFV | MP12 | MDCK | >100 |  |
|  |  | VeroE6 | 2.2 ± 0.23 |  | TPMV | VRC-66412 | VeroE6 | >100 |  |
| HCoV | 229E | MRC5 | 0.94 ± 0.16 |  | IAV H5N1 | A/Hong Kong/483/97 | A549 | >100 |  |
| HCoV | OC43 | MRC5 | 4.8 ± 0.25 |  | IAV H7N9 | A/Anhui/1/2013 | MA104/<br>TMPRSS2 | >100 |  |
| qPCR Assay Results |  |  |  |  |  |  |  |  |  |
| Virus | Strain | Cell | EC<sub>50</sub> (μM) <sup>d</sup> | EC<sub>90</sub> (μM) <sup>d</sup> | Virus | Strain | Cell | EC<sub>50</sub> (μM) <sup>d</sup> | EC<sub>90</sub> (μM) <sup>d</sup> |
| DENV2 | D2/hu/INDIA/09-74 | VeroE6 | 2.4 | 7.4 | SARS-CoV-2 | WK-521 (Ancestral) | VeroE6/<br>TMPRSS2 | 1.0 | 4.3 |
| ZIKV | MR766 | VeroE6 | 3.7 | 22 |  | QK002 (Alpha) | VeroE6/<br>TMPRSS2 | 0.64 | 2.3 |
| YFV | 17D-204 | VeroE6 | 0.36 | 11 |  | TY8-612 (Beta) | VeroE6/<br>TMPRSS2 | 1.7 | 7.6 |
| JEV | Beijing-1 | VeroE6 | 7.5 | 29 |  | TY7-501 (Gamma) | VeroE6/<br>TMPRSS2 | 1.3 | 3.6 |
| WNV | NY99 | VeroE6 | 0.60 | 10 |  | TY11-927 (Delta) | VeroE6/<br>TMPRSS2 | 0.25 | 2.1 |
| CHIKV | SL10571 | VeroE6 | 1.7 | 13 |  | TY38-873 (Omicron) | VeroE6/<br>TMPRSS2 | 0.69 | 4.3 |
| HCoV | 229E | MRC5 | 1.3 | 6.2 | RABV | HEP | BHK-21 | >40 | >40 |
| HCoV | OC43 | VeroE6 | 1.1 | 4.2 | RVFV | MP12 | VeroE6 | >40 | >40 |
| SARS-CoV | Hanoi | VeroE6 | 1.2 | 4.5 | HSV-1 | F | VeroE6 | >40 | >40 |
| MERS-CoV | EMC2012 | VeroE6/<br>TMPRSS2 | 2.3 | 8.2 |  |  |  |  |  |
| Resazurin Assay Results |  |  |  |  |  |  |  |  |  |
| Virus | Strain | Cell | EC<sub>50</sub> (μM) <sup>b</sup> | Fold change <sup>e</sup> |  |  |  |  |  |
| rgDENV2 | Wild type | BHK-21 | 0.86 ± 0.12 |  |  |  |  |  |  |
| rgDENV2 | NS5-G605V | BHK-21 | 5.1 ± 0.69 | 6.1 ± 0.51 |  |  |  |  |  |

Antiviral assays were carried out as described in Supplementary Table. EC<sub>50</sub>: 50% effective concentration. EC<sub>90</sub>: 90% effective concentration. CC<sub>50</sub>: 50% cytotoxic concentration.

Next, we performed cell-based antiviral assays (MTT assay or resazurin assay) using several RNA viruses to assess whether the antiviral activity of s2U was effective across a broad spectrum of viruses. s2U exhibited sub-micromolar to micromolar antiviral activity against ssRNA+ viruses, including viruses in the *Flaviviridae, Togaviridae* and *Coronaviridae* families (Table 1). The cell-based assay revealed that the 50% cytotoxic concentration (CC_50_) value of s2U was high (>400 μM for BHK-21, Vero E6 and MRC5) (Extended Data Fig. 1b). On the other hand, s2U did not show antiviral activity against negative-sense single-stranded RNA (ssRNA-) viruses, including rabies virus (RABV), Rift Valley fever virus (RVFV) and IFV (Table 1).

Real-time quantitative reverse transcription PCR (qRT-PCR) analysis and immunofluorescence assays confirmed that s2U inhibited the replication of the ssRNA+ viruses (DENV2, ZIKV, YFV, JEV, WNV, CHIKV, human coronavirus [HCoV]-229E, HCoV-OC43, SARS-CoV and Middle East respiratory syndrome coronavirus [MERS-CoV]) in a dose-dependent manner (Fig. 1b-1k, Table 1, Extended Data Fig. 2). However, s2U did not inhibit viral genome replication of ssRNA-viruses, such as RABV and RVFV, and the DNA virus herpes simplex virus-1 (HSV-1) (Fig. 1n-1p, Table 1). Furthermore, we observed dose-dependent inhibition of viral protein expression in DENV2-, CHIKV- and HCoV-OC43-infected cells (Fig. 1m). Notably, s2U also inhibited viral RNA replication in the SARS-CoV-2 ancestral strain and each variant of concern (VOC), including the Delta (B.1.617.2 lineage) and Omicron (BA.1 lineage) strains, in a dose-dependent manner (Fig. 1l, Table 1, Extended Data Fig. 2). Viral protein expression in SARS-CoV-2-infected cells was inhibited by s2U in a dose-dependent manner (Fig. 1m).

### s2U blocks RNA synthesis by stalling of viral RdRp

We performed antiviral assays simultaneously treated with various doses of the four ribonucleosides (adenosine, guanosine, uridine, and cytidine) to confirm whether s2U acts as a nucleoside analogue. The antiviral activity of s2U was reduced following the addition of an excess of exogenous pyrimidine ribonucleosides (uridine and cytidine) to the infected cells but not purine ribonucleosides (adenosine and guanosine) (Fig. 2a, Extended Data Fig. 3a), suggesting that s2U inhibits viral RNA replication by acting as a uridine decoy.

**Fig. 2.**
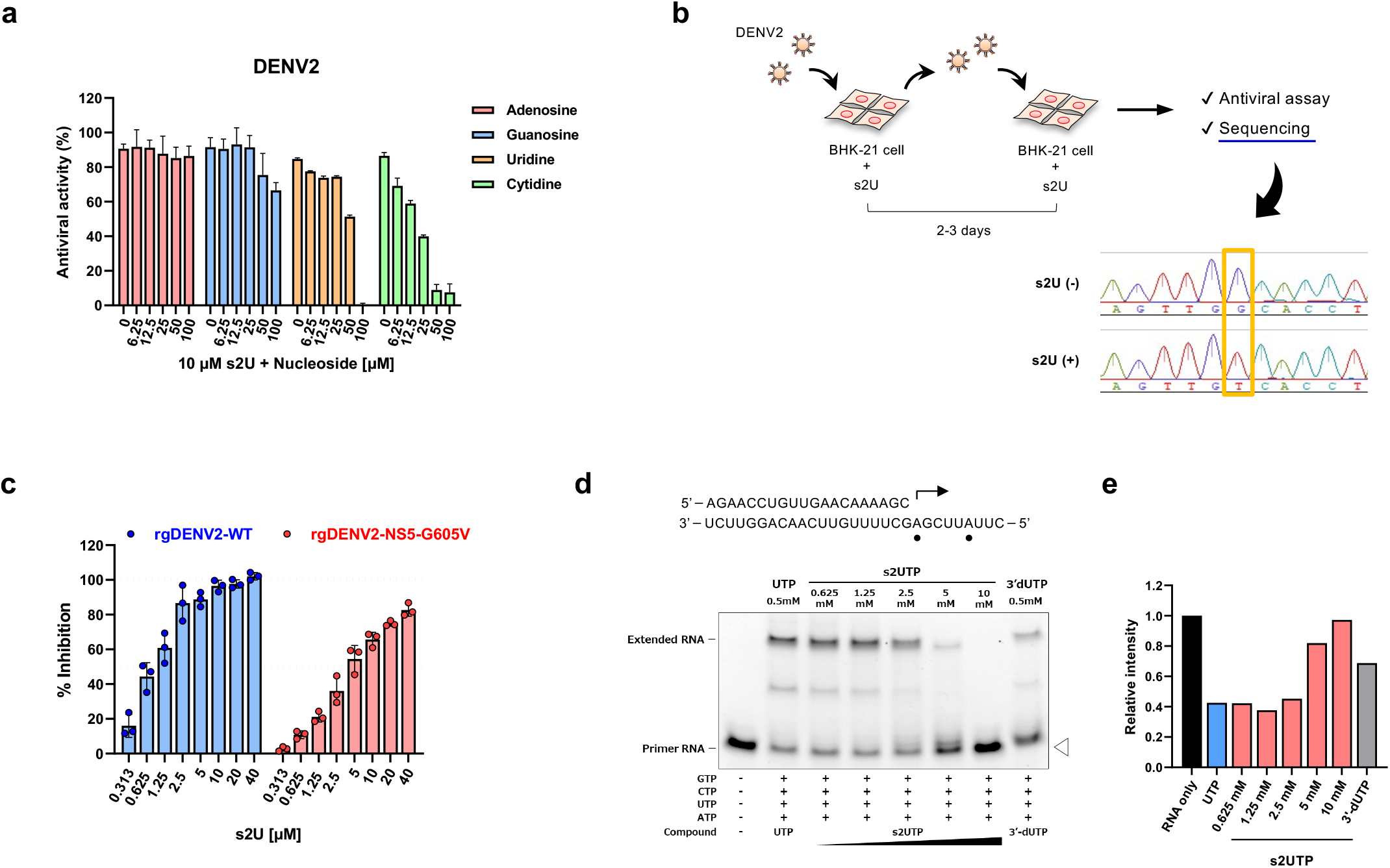
Molecular target and mechanism of action of s2U. **a**, Ribonucleotide competition of DENV2 inhibition by s2U. DENV2 (multiplicity of infection [MOI] = 0.01)-infected BHK-21 cells were treated with 10 μM of s2U and serial dilutions of exogenous nucleosides. A resazurin reduction assay was performed at 4 days post infection (dpi). **b**, Schematic of the experimental design for drug-escape mutant selection and sequence result. BHK-21 cells were infected with DENV2 in the presence of s2U. The passage of infected cells or culture supernatant was performed every 2–3 days. Base substitution was detected using Sanger sequencing. **c**, Effect of s2U resistance mutation on anti-DENV2 activity of s2U. BHK-21 cells were infected with rgDENV2-WT or rgDENV2-NS5-G605V (MOI = 0.1) containing a serially diluted compound. A resazurin reduction assay was performed at 4 dpi. **d**, Analysis of viral RdRp stalling by s2U 5’-triphosphate (s2UTP). Denaturing polyacrylamide gel electrophoresis fraction of RNA transcripts produced through primer extension by ZIKV RdRp in the presence of the indicated nucleotides. The RNA primer/template sequence used in this assay is indicated at the top (small black circles indicate the incorporation sites of UTP). **e**, Relative band intensities of fluorescently labeled RNA primers. Relative fluorescence intensities of each RNA primer (white arrowhead in Fig. 2d) were normalized by the RNA sample without UTP or s2UTP (black bar, RNA only). Anti-DENV2 activities (%; **a, c**) are expressed relative to the values for the DMSO-treated, infected samples and non-infected samples. Data are presented as mean values, and error bars indicate SD.

To identify the molecular target of s2U, we selected a drug-escape mutant by passaging DENV2 in the presence of gradually increasing s2U concentrations. We observed an almost complete decrease in s2U susceptibility at passage 19 and identified a single base substitution (G1814T) within the viral non-structural protein 5 (NS5) in this escape mutant virus (Fig. 2b). This single mutation resulted in a single amino acid substitution (G605V) in the RdRp coding region, which was not observed in the in-parallel-passaged viruses in the absence of s2U.

Next, we constructed recombinant DENV2 possessing NS5-G605V (rgDENV2-WT and rgDENV2-NS5-G605V) using a reverse genetics system^25^ to evaluate replication fitness and s2U resistance caused by the G605V mutation. The mutation did not affect viral replication in different cell lines (Extended Data Fig. 3b, 3c). Although the G605V mutation slightly affected the sensitivity of ribavirin and favipiravir in the antiviral assay (2.3- and 2.2-fold decrease, respectively) (Extended Data Table 1), it reduced the sensitivity to s2U by 6.1-fold compared with recombinant rgDENV2-WT (Table 1, Fig. 2c).

Nucleoside analogues can sometimes serve as substrates for mitochondrial RNA polymerase (POLRMT), resulting in mitochondrial toxicity and side effects^26^. To assess the impact of s2U on POLRMT and RNA polymerase II (RNA pol II) activity, we performed mitochondrial protein synthesis assays using an In-Cell ELISA method^26^. s2U did not affect the steady-state levels of mitochondrial-encoded (cytochrome *c* oxidase I [COX-I]) and nuclear-encoded (succinate dehydrogenase [SDH]-A) proteins, suggesting that POLRMT and RNA pol II activities were unaffected (Extended Data Fig. 4).

To characterize the molecular mechanism of action of s2U, we performed *in vitro* primer extension assay^27^ using ZIKV NS5 protein and s2U 5’-triphosphate (s2UTP). Although UTP was incorporated into the RNA template and extended this, the incorporation of s2UTP instead of UTP might likely block RNA extension in the presence of the next correct ribonucleotide (Fig. 2d, 2e). These results suggest that s2UTP acts on viral RdRp and inhibits viral RNA synthesis.

### *In vivo* efficacy of s2U against DENV2

Subsequently, we employed an animal model to evaluate the *in vivo* antiviral activity of s2U. A mouse-adapted DENV2 model has been established using AG129 (interferon [IFN]-α/β and IFN-g receptor-deficient 129/Sv) mice^28^, and this model has been utilized to develop of anti-DENV drugs and vaccines^29–31^. Thus, we established a mouse-adapted DENV2 strain (DENV2 AG-P10) based on this model^28^ to evaluate the *in vivo* efficacy of the compound. This mouse-adapted DENV2 strain had two amino acid substitutions (NS4B-A119T and NS5-E802Q) that developed during viral passages and demonstrated higher pathogenicity in AG129 mice compared to the parental clinical isolate (Extended Data Fig. 5).

AG129 mice were intraperitoneally inoculated with DENV2 AG-P10 and treated twice daily with s2U (50 or 150 mg/kg of body weight [BW]) by oral gavage starting immediately after infection (Fig. 3a). After 5 consecutive days of treatment, the survival rate of virus-inoculated AG129 mice was significantly increased in the mice that received the s2U treatment (median survival: 12.5 days at 50 mg/kg and >16 days at 150 mg/kg) compared with the vehicle-treated mice (median survival: 9 days) (Fig. 3b). A dose-dependent decrease in viral RNA load at 3 days post infection (dpi) was also observed in serum, spleen, kidney and liver samples (Fig. 3c–3f). These data suggest that s2U protects against DENV2-induced mortality by decreasing viral propagation *in vivo* in AG129 mice.

**Fig. 3.**
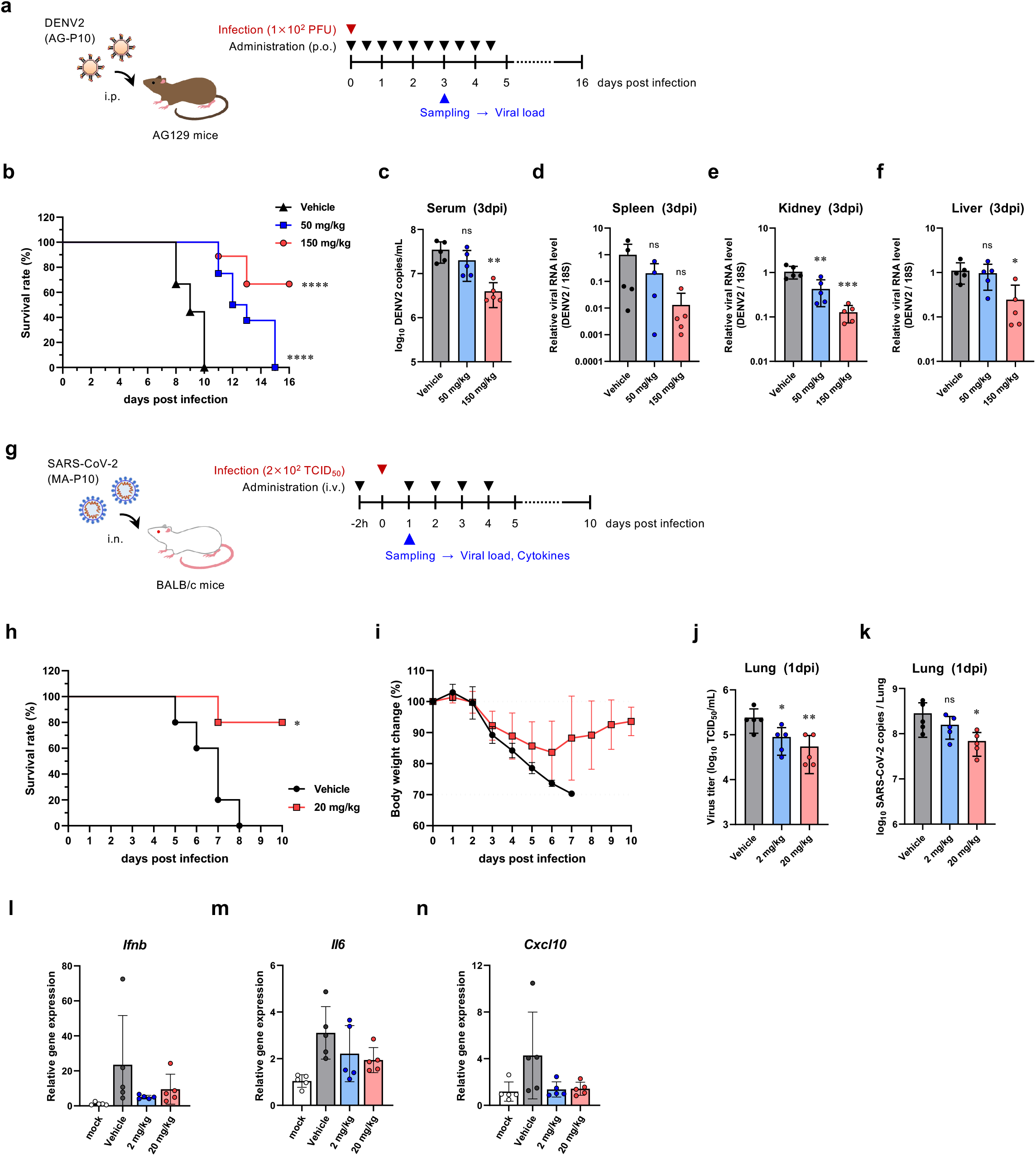
*In vivo* efficacy of s2U in the DENV2 and SARS-CoV-2 mouse model. **a**, Schematic representation of the survival and viremia studies using AG129 mice and strain DENV2 AG-P10. **b**, Effect of s2U on survival of DENV2 AG-P10-infected (1 × 10^2^ plaque forming units [PFU]) mice orally treated twice daily with s2U (50 or 150 mg/kg) compared with vehicle-treated mice. Treatment started immediately after infection. Data are from two independent studies with 9 (in total, vehicle and 150 mg/kg) or 8 (in total, 50 mg/kg) mice per group. **c, d**, Effect of s2U on viremia at 3 dpi in mice treated twice daily with s2U (50 or 150 mg/kg) compared with vehicle-treated mice (n = 5 per group). Viral RNA copies/mL of serum samples (**c**) or the relative viral RNA level (DENV2 copies/18S copies) of spleen (**d**), kidney (**e**) and liver (**f**) samples were quantified using qRT-PCR. **g**, Schematic representation of the survival and viremia studies using BALB/c mice and SARS-CoV-2 MA-P10. **h, i**, Effect of s2U on survival and body weight change in SARS-CoV-2 MA-P10-infected (2 × 10^2^ 50% tissue culture infection dose [TCID_50_]) mice intravenously administered 20 mg/kg s2U daily compared with vehicle-treated mice (n = 5 per group). Treatment started 2 h before infection. Survival (**h**) and body weight (**i**) of the mice were monitored daily. **j, k**, Effect of s2U on viremia at 1 dpi in mice intravenously administered 2 or 20 mg/kg s2U daily compared with vehicle-treated mice (n = 5 per group). Virus titers in lung samples (**j**) were quantified by a standard TCID_50_ assay using VeroE6/TMPRSS2 cells. Viral RNA copies/mL in lung samples (**k**) were quantified using qRT-PCR. **l, m, n**, Relative *Ifnb* (**l**), *Il6* (**m**) and *Cxcl10* (**n**) gene expression profiles in lungs from mice at 1 dpi with SARS-CoV-2. Cytokine RNA levels were determined relative to *18S rRNA* transcripts. Data are presented as mean values, and error bars indicate SD. Statistically significant differences between the s2U-treated and vehicle-treated groups were determined using a Log-rank (Mantel–Cox) test (**b**, **h**) and one-way ANOVA followed by Dunnett’s multiple comparisons test (**c**, **d**, **e**, **f**, **j**, **k**); * *p* < 0.01, ** *p* < 0.005, *** *p* < 0.0005 and **** *p* < 0.0001.

### *In vivo* efficacy of s2U against SARS-CoV-2

SARS-CoV-2 does not show infectivity and pathogenicity to normal mice. Thus, human angiotensin-converting enzyme 2 transgenic mice are widely used as a small-animal model of SARS-CoV-2 infection^32,33^. A mouse-adapted SARS-CoV-2 model was recently established, which has promoted research on SARS-CoV-2 pathogenicity and the development of drugs and vaccines against COVID-19^34,35^. Therefore, to evaluate the *in vivo* efficacy of s2U, we also established a mouse-adapted SARS-CoV-2 strain (SARS-CoV-2 MA-P10) by following previous reports^36,37^ (Extended Data Fig. 6a, 6b). The resultant mouse-adapted virus harbors a G498H substitution in the spike protein and can efficiently replicate in normal mice (Extended Data Fig. 6c–6g). These results were consistent with reports of other mouse models^36–39^.

The effect of s2U on virus-induced mortality was assessed by daily intravenous administration (20 mg/kg) for 5 consecutive days starting 2 h before infection (Fig. 3g). BALB/c mice (30–50 weeks old) were intranasally infected and then monitored for a maximum of 10 days. We observed a statistically significant increase in survival rate (80% survival) after 5 days of s2U treatment compared with the vehicle-treated mice (Fig. 3h). The peak BW loss in mice treated with s2U was approximately 17%, and the BW was recovered following the administration of s2U, while the BW of vehicle-treated mice continued to decrease daily (Fig. 3i).

The mouse-adapted SARS-CoV-2 replicates rapidly, and the viral load in the lungs peaks at 1 dpi (Extended Data Fig. 6d, 6e)^37^. Therefore, we next assessed the effect of s2U on the viral load at 1 dpi following intravenous administration (20 mg/kg) to 5-week-old infected BALB/c mice. A dose-dependent decrease in viral titer and viral RNA load was observed in the lungs of s2U-treated mice compared with vehicle-treated mice (Fig. 3j, 3k). We also observed decreased viral titers in the lungs following twice-daily oral administration of s2U (300 mg/kg) immediately after infection (Extended Data Fig. 7a, 7b). Finally, we detected lower levels of several pro-inflammatory cytokines (IFN-β, IL-6 and CXCL10) in lung homogenates prepared from mice treated with s2U compared with those of vehicle-treated mice (Fig. 3l–3n). These data suggest that s2U protects against SARS-CoV-2-induced lung inflammation and mortality by decreasing viral propagation.

## Discussion

This study identified the antiviral ribonucleoside analogue s2U as a strong inhibitor of viral replication of different clinically relevant ssRNA+ virus families, including *Flaviviridae*, *Togaviridae* and *Coronaviridae,* without significant cytotoxicity. Additionally, Alam and colleagues reported that s2U inhibits the replication of murine norovirus-1 (MNV-1)^40^, which belongs to the *Caliciviridae* family. Collectively, our findings indicate that s2U inhibits the replication of ssRNA+ viruses but not ssRNA– and DNA viruses. These findings suggest that the broad-spectrum antiviral activity of s2U against ssRNA+ viruses is due to a unique mechanism of action.

Our studies demonstrated an s2U-escape mutant with a G605V substitution in the NS5 protein of DENV2 D2/hu/INDIA/09-74 strain, which is located at motif B of DENV2 RdRp (corresponding to Gly604 described in ref. 41). Moreover, s2UTP inhibited RNA extension catalyzed by the ZIKV RdRp. s2UTP has been reported to be incorporated into the RNA template of feline calicivirus and human norovirus by ProPol (a precursor comprised of both the proteinase and polymerase), resulting in the inhibition of norovirus and calicivirus polymerase activity, respectively^42^. Another study provided structural evidence of s2U binding to MNV-1 RdRp and showed that s2U interacted with Thr309 and other amino acids that form the active site pocket^40^. Thr309 is positioned in a motif B of MNV-1 RdRp, which corresponds to the same region as Gly605 of DENV2 RdRp^41^ and Ala688 of SARS-CoV-2 RdRp^ref^. These findings suggest that the molecular target of s2U is the active site of ssRNA+ viral RdRp. Sofosbuvir (SOF)^43^ and 4’-fluorouridine (4’-FIU)^44^ are potent uridine analogues against hepatitis C virus and RSV, respectively. 4’-FIU also inhibits SARS-CoV-2 replication *in vitro* and *in vivo*^44^. SOF and 4’-FIU 5’-triphosphates (TP) are incorporated into the RNA template by mimicking UTP, inducing stalling of viral RdRp^44,45^. SOF-TP is also incorporated into the RNA template by ZIKV RdRp, resulting in inhibition of viral RNA synthesis^27^. Our study demonstrated s2U as another uridine analogue, and s2UTP induced stalling of ZIKV RdRp. These findings suggest that s2U inhibits viral RNA synthesis by blocking the activity of the viral RdRp, while further investigation including comprehensive *in vitro* RNA extension assay and s2U complex structure of SARS-CoV-2 RdRp is necessary to conclude the detailed mechanism of action of s2U.

Once-daily intravenous and twice-daily oral administration of s2U to mice significantly reduced the viral load of SARS-CoV-2 and DENV2, respectively. Moreover, continuous administration of s2U to mice significantly increased the survival rate of both SARS-CoV-2- and DENV2-infected mice. In our pharmacokinetic analysis, simulation of the repeated dose concentration-time profile revealed that once-daily intravenous and twice-daily oral administration of s2U maintains the validated concentration of more than 50% effective concentration (EC_50_) or 90% effective concentration (EC_90_) against DENV2 and SARS-CoV-2 (Extended Data Fig. 8). s2U exhibited broad anti-coronaviral efficacy with equally strong activity against SARS-CoV-2 VOCs *in vitro,* except for Omicron variant, which has somehow distinct feature and whose infectious activity is inhibited but with 2-log10 fold reduction. Thus, we can conclude that the s2U will retain its efficacy against future variants of SARS-CoV-2 which may have substantial resistance to spike-targeting antibody therapeutics or vaccines. Furthermore, s2U also exhibited broad anti-flavivirus efficacy with equally strong activity *in vitro*. These findings suggest the high potential of s2U as an efficacious broad-spectrum oral and/or intravenous agent against ssRNA+ viruses, making it a promising therapeutic option for COVID-19, Dengue and other diseases caused by ssRNA+ viruses.

In conclusion, we demonstrated that s2U exhibits strong, safe, broad-spectrum antiviral activity against ssRNA+ viruses by inhibiting viral RdRp activity. Viral RdRp represents a promising target for the development of broad-spectrum inhibitors because it is functionally and structurally conserved across among a wide range of viruses. Thus, the use of s2U (and s2U-derivatives) may be extended for the development of new drugs against related or novel viruses, and these compounds may possibly contribute to our progress in the area of pandemic preparedness.

## Methods

### Cell lines

BHK-21 (ATCC, CCL-10), VeroE6 (ATCC, CRL-1586), MA104 (RIKEN BRC, RCB0994), 293T (RIKEN BRC, RCB2202), HepG2 (ATCC, HB-8065), A549 (RIKEN BRC, RCB0098), KB (ATCC, CCL17), MDBK (ATCC, CCL-22) and MDCK (ATCC, CCL-34) cells were maintained in high-glucose Dulbecco’s modified Eagle’s medium (DMEM, Gibco) supplemented with 10% fetal bovine serum (FBS, Gibco) and penicillin–streptomycin (P/S, Wako) at 37°C. MRC5 (ATCC, CCL-171) cells were also maintained in Minimum Essential Medium GlutaMAX Supplement (Gibco) supplemented with 10% FBS, nonessential amino acids (NEAA, Wako), sodium pyruvate (Wako) and P/S at 37°C. THP-1 (ATCC, TIB-202) and MOLT4 (JCRB, JCRB9031) cells were maintained in Roswell Park Memorial Institute 1640 (RPMI-1640, Gibco) supplemented with 10% FBS and P/S at 37°C. C6/36 (ATCC, CRL-1660) cells were maintained in Minimum Essential Medium (MEM, Nissui) supplemented with 10% FBS, NEAA and L-Alanyl-L-glutamine (Wako) at 28°C.

### Generation of TMPRSS2-expressing cells

VeroE6 cells stably expressing human TMPRSS2 (VeroE6/TMPRSS2) were generated by lentiviral transduction with CSII-CMV-TMPRSS2-IRES2-Bsd and blasticidin-based selection as described in previous report^46^. MA104 cells stably expressing human TMPRSS2 (MA104/TMPRSS2) were also generated as same method. For the lentiviral vector preparation, 293T cells were co-transfected with the aforementioned lentiviral vector plasmid and Lentiviral High Titer Packaging Mix (Takara Bio).

### Viruses

DENV1 (D1/hu/PHL/10–07 strain), DENV2 (D2/hu/INDIA/09–74 strain, GenBank: LC367234), DENV3 (D3/hu/Thailand/00–40 strain), DENV4 (D4/hu/Solomon/09–11 strain), ZIKV (MR766 strain, GenBank: LC002520), YFV (17D–204 strain), JEV (Beijing–1 strain) and WNV (NY99 strain) were propagated as described in previous report^25^.

CHIKV strain SL10571 was propagated as described in previous report^47^.

Human coronavirus strain OC43 (HCoV-OC43) and 229E (HCoV-229E) were purchased from ATCC (VR-1558 and VR-740, respectively) and amplified as described in previous report^48^.

SARS-CoV strain Hanoi was kindly provided by Nagasaki University and was amplified in VeroE6 cells^49^.

MERS-CoV strain EMC2012 was kindly provided by Erasmus University Medical Center and amplified on VeroE6/TMPRSS2 cells^50^.

SARS-CoV-2 strain WK-521 (Ancestral, Pango Lineage: A, GISAID: EPI_ISL_408667), QK002 (Alpha, Pango Lineage: B.1.1.7, GISAID: EPI_ISL_768526), TY8-612 (Beta, Pango Lineage: B.1.351, GISAID: EPI_ISL_1123289), TY7-501 (Gamma, Pango Lineage: P.1, GISAID: EPI_ISL_833366), TY11-927 (Delta, Pango Lineage: B.1.617.2, GISAID: EPI_ISL_2158617), TY38-873 (Omicron, Pango Lineage: BA.1, GISAID: EPI_ISL_7418017), a clinical isolate from a patient with COVID-19, were kindly provided by National Institute of Infectious Diseases (NIID); the original stock of these virus strains were prepared by inoculation of VeroE6/TMPRSS2 cells.

RABV high egg passage Flury (HEP) strain was propagated as described in previous report^51^.

La Cross virus (LACV) was purchased from ATCC (VR-1834) and amplified on BHK-21 cells.

Leopards Hill virus (LPHV, 11SB17 strain) was amplified in KB cells as previously described^52^.

Lymphocytic choriomeningitis virus (LCMV, Armstrong strain) and Junin virus (JUNV, Candid #1 strain) were kindly provided by NIID and was amplified in BHK-21 cells.

Severe fever with thrombocytopenia syndrome virus (SFTSV, ArtLN/2017 strain) was propagated as described in previous report^53^.

Recombinant Rift Valley fever virus (RVFV, MP12 strain) was rescued as described previously^54^ and was amplified in BHK-21 cells.

Thottopalayam thottimvirus (TPMV, VRC-66412 strain) was kindly provided by Hokkaido University and was amplified in VeroE6 cells^55^.

Avian influenza A virus H5N1 (IAV-H5N1, A/Hong Kong/483/97 strain) was kindly provided by University of Hong Kong and was propagated in embryonated chicken eggs and harvested from virus-containing allantoic fluids.

Avian influenza A virus H7N9 (IAV-H7N9, A/Anhui/1/2013 strain) was kindly provided by NIID and was propagated in embryonated chicken eggs and harvested from virus-containing allantoic fluids.

Herpes simplex virus (HSV-1, F strain) was kindly provided by NIID and was amplified in VeroE6 cells.

WNV, CHIKV, SARS-CoV, MERS-CoV, SARS-CoV-2, SFTSV, RVFV, IAV-H5N1 and IAV-H7N9 were propagated in a biosafety level-3 (BSL-3) facility at the International Institute for Zoonosis Control, Hokkaido University.

### Compounds

All compounds that were screened, including 2-thiouridine, were synthesized at the Faculty of Pharmaceutical Sciences, Hokkaido University and their chemical identity and purity were determined by high-performance liquid chromatography and mass spectrometry analysis. Ribavirin, Favipiravir, GS-5734 (Remdesivir), GS-441524, 2-thiouridine and Chloramphenicol were purchased from Sigma-Aldrich, PharmaBlock Sciences, Inc., MedChemExpress, Carbosynth Limited, Cayman Chemical Company and Calbiochem, respectively. For *in vitro* studies, all compounds were solubilized in 100% dimethyl sulfoxide (DMSO; Sigma-Aldrich) and were diluted in 2% FBS/MEM. For *in vivo* studies, 2-Thiouridine was dissolved in DMSO and diluted with 0.5% methylcellulose (MC) aqueous solution to prepare 50 or 150 mg/mL solutions for oral administration. 2-Thiouridine was also dissolved in OTSUKA NORMAL SALINE (Otsuka Pharmaceutical Co., Ltd.) for intravenous administration.

### Cell-based Antiviral and Cytotoxicity Assays

The MTT (3-[4,5-dimethyl-2-thiazolyl]-2,5-diphenyl-2H-tetrazolium bromide) and resazurin reduction assays were carried out as previously described^25,48,56^. These were performed to calculate cell viability following viral induced cytopathic effect (CPE) and cytotoxicity. Assay conditions for all viruses are as described in Supplementary Table 1. Cells and viruses were incubated in 96-well plates with the 2-fold serially diluted compound (n = 2) in all assays. The EC_50_ value was defined in GraphPad Prism version 8.4.3 (GraphPad Software) with a variable slope (four parameters). Non-infected cells were used as a control for 100% inhibition, whereas for infected cells, DMSO alone was used as a control for 0% inhibition. The CC_50_ value for each cell line was also measured using the same method. Cell-free samples were used as 100% cytotoxicity control and DMSO-treated cells were used as 0% cytotoxicity control.

### Quantification of viral RNA with real-time quantitative reverse transcription PCR (qRT–PCR)

Assay conditions for all viruses are as described in Supplementary Table 1. Briefly, cells were seeded onto 48-well plates the previous day and infected with the virus containing the serially diluted compound. After a certain period of time, total RNA was isolated with PureLink RNA Mini Kit (Ambion; Thermo Fischer Scientific). Viral RNA from all samples was quantified using real-time RT-PCR analysis with EXPRESS One-step Superscript qRT-PCR kit (Invitrogen; Thermo Fischer Scientific) and QuantStudio 7 Flex Real-Time PCR system (Applied Biosystems; Thermo Fischer Scientific). The primers and probe sequences were designed in previous reports and are described in Supplementary Table 2, with primers and probe for *ACTB* (Hs01060665_g1, Applied Biosystems) and *18S rRNA* (Hs99999901_s1, Applied Biosystems) transcripts used as internal controls. The EC_90_ value was defined in GraphPad Prism version 8.4.3 with a variable slope (Find ECanything, F=90).

### Quantification of viral DNA with real-time quantitative PCR (qPCR)

VeroE6 cells were seeded the previous day and infected with HSV-1 at a multiplicity of infection (MOI) of 0.1 containing the serially diluted compound. At 24 hours post-infection (hpi), total DNA was isolated with DNeasy Blood & Tissue Kit (Qiagen). Viral DNA from samples was quantified using real-time PCR analysis with TaqMan Fast Advanced Master Mix (Applied Biosystems) and QuantStudio 7 Flex Real-Time PCR system. The primers and probe sequences were designed in previous reports and are described in Supplementary Table 2, with primers and probe for *ACTB* transcripts used as internal controls.

### Indirect immunofluorescence assay (IFA)

Assay conditions for all viruses are as described in Supplementary Table 1. Briefly, cells were seeded onto 48-well plates the previous day and infected with the virus containing the serially diluted compound. After a certain period of time, cells were fixed with the buffered formalin (Masked Form A, Japan Tanner Co.), permeabilized with 0.5% Triton X-100 in PBS or ice-cold methanol. Cells were then stained with the 4G2 (supernatant from D1-4G2-4-15 cells, ATCC HB-122), Anti-Chikungunya virus E1 mAb, clone 1B6 (MAB12424, Abnova), anti-coronavirus antibody (MAB9013, Merck Millipore), SARS-CoV-2 nucleocapsid antibody (HL344; GTX635679, GeneTex) and Alexa Fluor Plus 488-conjugated anti-mouse or anti-rabbit IgG antibody (Invitrogen). Cell nuclei were counterstained with Hoechst 33342 (Molecular Probes). Cells were then evaluated using fluorescence microscopy (IX73, Olympus). Images were processed with cellSens Standard 1.16 (Olympus).

### Ribonucleotide competition of DENV2 and HCoV inhibition

BHK-21 or MRC5 cells were infected with DENV2 (MOI = 0.01) or HCoV-229E (MOI = 0.005), assay media was supplemented with s2U at 10 or 15 μM alone or in combination with 6.25 to 100 μM exogenous ribonucleosides (Adenosine, Guanosine, Uridine, Cytidine, Sigma-Aldrich). Resazurin reduction assay was carried out at 96- or 72-hours post-infection, respectively. Antiviral activities (%) are expressed relative to the values for the DMSO-treated, infected samples and non-infected samples.

### Isolation of drug-escape mutant

BHK-21 cells seeded onto 12-well plates and were infected with DENV2 at an MOI of 0.01 containing the 2 μM or 6 μM of s2U. The passage of infected cells was performed every 2–3 days. When a CPE was observed, culture supernatants were transferred to non-infected BHK-21 cells in the presence of the compound. After continuous culture of the viruses for 42 days (passage 19), the viral RNA was isolated and amplified by RT-PCR using the PrimeScript II High Fidelity One Step RT-PCR Kit (Takara Bio) and specific primers. Viral genome was then examined by Sanger sequencing using specific primers and the 3500xL Genetic Analyzer (Life Technologies). The sequence was compared to that of wild-type virus.

### Construction of recombinant and mutant DENV2 infectious clones

Recombinant DENV2 (D2/hu/INDIA/09-74, rgDENV2-WT) was rescued and amplified as described previously^25^. A point mutation fragment was amplified using PCR-based site directed mutagenesis method and inserted into a plasmid containing the whole DENV2 genome. The T3 promoter and DENV2 genomic region of the plasmid was amplified by PCR using KOD One PCR Master Mix -Blue- (Toyobo), and DENV2 genomic RNA was synthesized using a mMessage mMachine T3 Kit (Thermo Fisher Scientific). The genomic RNA was transfected into BHK-21 cells with *Trans*IT-mRNA reagent (Mirus), and culture supernatants were collected when CPEs were observed.

### *In vitro* growth kinetics of drug-resistant mutant

VeroE6 cells seeded onto 24-well plates the previous day and were infected with rgDENV2 (rgDENV2-WT or rgDENV2-NS5-G605V) at an MOI of 0.01 for 1 h. After incubation, the unbound virus was removed, and new medium was added. At 24, 48, 72 and 96 hpi, total RNA was isolated with PureLink RNA Mini Kit. Viral RNA level was quantified by qRT-PCR analysis as described above with *ACTB* transcripts used as internal controls.

### Primer extension polymerase activity assay

The primer extension assay was performed according to a previous report^27,48^. For analysis of the competitive inhibition ability of s2UTP, 200 nM of recombinant ZIKV NS5 (40546-V08B, Sino Biological) and 50 nM RNA primer-template complexes (RNA P/T) were incubated 30°C for 15 min in reaction buffer [10 mM Tris-HCl (pH 7.5), 10 mM DTT, 5 mM MgCl_2_, 5% glycerol, 0.05% Triton-X 100, and 0.02 U/μL RNasein (Promega)], and then UTP analogues [500 μM UTP (Thermo Fischer Scientific), 500 μM 3’-dUTP (TriLink), or serially diluted s2UTP (0.625 to 10 mM)] were added and incubated at 30°C for 10 min. Primer extension reactions were initiated by the addition of ATP, GTP, CTP and UTP (100 μM each). The reactions were performed 30°C for 2 h and stopped by the addition of quenching buffer (7M Urea, 1X TBE buffer, and 50 mM EDTA). The quenched samples were denatured at 95°C for 5 min, and the primer extension products were separated on a 15% denaturing polyacrylamide gel (Invitrogen). After electrophoresis, the gels were scanned using an Amersham ImageQuant 800 Fluor system (Cytiva). The band intensities were analyzed by ImageQuant TL version 8.2.0 (Cytiva).

### Mitochondrial protein synthesis assay

HepG2 cells were seeded onto 96-well plates the previous day and treated with the 3-fold serially diluted compound (n = 2). At 5 dpi, one set of plates was fixed with 4% paraformaldehyde (Nacalai tesque) and determined the intracellular level of two mitochondrial proteins, the mitochondrial DNA-encoded cytochrome *c* oxidase I (COX-I) and nuclear DNA-encoded succinate dehydrogenase A (SDH-A) using the MitoBiogenesis In-Cell Enzyme-linked immunosorbent assay (ELISA) Kit (Colorimetric, Abcam, ab110217) following the manufacturer’s instructions. To monitor the cell viability as ATP level, the second set of plates was analyzed by adding CellTiter-Glo 2.0 Reagent (Promega, G9242/3) and measuring luminescence on a GloMax Discover System (Promega). The IC_50_ value was defined in GraphPad Prism version 8.4.3 with a variable slope (four parameters).

### Ethical statement

All the animal experiments were performed in accordance with the National University Corporation, Hokkaido University Regulations on Animal Experimentation. The protocol was reviewed and approved by the Institutional Animal Care and Use Committee of Hokkaido University (approval no. 18-0046, 18-0149 and 20-0060).

### Establishment of mouse-adapted DENV2 and SARS-CoV-2

AG129 mice (IFN-α/β and IFN-g receptors deficient 129/Sv mice) were purchased from Marshall BioResources and bred in-house under the specific pathogen-free (SPF) condition. To establish mouse-adapted DENV2, named DENV2 AG-P10 strain, virus passage in mouse was carried out according to a previous report^28^. Briefly, 7-weeks-old female AG129 mice were inoculated intraperitoneally with 100 μL of 4×10^5^ PFU/mouse of DENV2 D2/hu/INDIA/09-74 strain. On day 3 after infection, the infected mice were euthanized under deep anesthesia by isoflurane inhalation, and serum was collected. Virus in the serum was amplified in C6/36 cells and then intraperitoneally injected into another AG129 mice. This adaptation process was performed a total of 10 times.

To establish mouse-adapted SARS-CoV-2, named SARS-CoV-2 MA-P10 strain, virus passage in mouse was carried out according to a previous report^36,37^. Briefly, specific pathogen-free (SPF), 30–45-week-old female BALB/c mice (BALB/cAJcl, CLEA Japan) were inoculated intranasally with 50 μL of 1×10^5^ TCID_50_/mouse of SARS-CoV-2 WK-521 strain under anesthesia. On day 3 after infection, the infected mice were euthanized, and whole lung tissues were harvested and homogenized in DMEM supplemented with 10% FBS and P/S with TissueRuptor (Qiagen). Virus in the supernatant of lung homogenate was intranasally injected into another BALB/c mice. This adaptation process was performed a total of 10 times.

Viral RNA was extracted and purified using QIAamp Viral RNA Mini Kit (Qiagen) according to the manufacturer’s instructions. Next-generation sequencing (NGS) was conducted on an iSeq 100 System (Illumina, Inc.) and the sequences were analyzed using CLC Genomics Workbench ver. 21.0.3 software (CLC bio, Qiagen).

### Determination of antiviral activity *in vivo*

In DENV2 infection model, SPF, sex matched 7-week-old AG129 mice were inoculated intraperitoneally with 100 μL of 2×10^2^ PFU/mouse of DENV2 AG-P10 strain. After inoculation, mice were treated with s2U twice daily by oral administration (50 or 150 mg per kg) for 5 consecutive days. Mice were monitored daily; body weight was determined daily. To evaluate the viral load, the infected mice were euthanized at 3 dpi, and serum, spleen, kidney and liver were collected and homogenized in PBS with TissueRuptor. The serum samples were subjected to standard plaque assay using BHK-21 cells for virus titration.

In SARS-CoV-2 infection model, SPF, 5-week-old (for viral load) or 30–50-week-old (for survival study) female BALB/c mice were inoculated intranasally with 50 μL of 2×10^2^ TCID_50_/mouse of SARS-CoV-2 MA-P10 strain under anesthesia. Mice were treated with s2U by intravenous (2 or 20 mg/kg, once daily) or oral (300 mg/kg, twice daily) administration. Treatment was initiated 2 h before inoculation and continued for 5 consecutive days for survival study. On day 1 after inoculation, the infected mice were euthanized, and whole lung tissues were harvested and homogenized in PBS with TissueRuptor. The homogenates were centrifuged for 10 min at 3,000 rpm to pellet tissue debris and the supernatants were subjected to standard TCID_50_ assay using VeroE6/TMPRSS2 cells for virus titration.

Viral RNA isolation from serum or tissue sample were performed using QIAamp Viral RNA Mini Kit (Qiagen) or PureLink RNA Mini Kit, respectively. Viral RNA level was quantified by qRT-PCR analysis as described above with *18S rRNA* transcripts used as internal controls.

### *In vitro* ADME assay

Solubility assay: The Japanese Pharmacopeia (JP) 1st fluid (pH 1.2) or JP 2nd fluid (pH 6.8) for dissolution testing was used for solubility measurements. A test solution of test compound was prepared by diluting 10 mM DMSO stock solution 2 μL:165 μL in JP1st or 2nd fluid and mixed at 37°C for 4 h by rotation at 1,000 rpm. After loading the mixed solution into 96-well MultiScreen Filter Plates (product number MSHVN4510, 0.45 μm hydrophilic PVDF membrane, Millipore), filtration was performed by centrifugation. The filtrates were mixed with acetonitrile and analyzed by HPLC-UV (254 nm). Solubility was calculated by comparing the peak area of the filtrate mixture with that of a 100 μM standard solution. When the peak area of the filtrate mixture was larger than the peak area of the standard solution, it was described as >100 μM.

PAMPA assay to determine the passive membrane diffusion rates: A Corning Gentest Pre-coated PAMPA Plate System was used in the PAMPA permeability test. The acceptor plate was prepared by adding 200 μL of 5% DMSO/0.1 M phosphate buffer (pH 7.4) to each well, and then 300 μL of 100 μM test compounds in 5% DMSO/0.1 M phosphate buffer (pH 6.4) was added to the donor wells. The acceptor plate was then placed on top of the donor plate and incubated at 37°C without agitation for 4 h. At the end of the incubation, the plates were separated and the solutions from each well of both the acceptor plate and the donor plate were transferred to 96-well plates and mixed with acetonitrile and water. The final concentrations of compounds in both the donor wells and acceptor wells, as well as the concentrations of the initial donor solutions, were analyzed by liquid chromatography tandem mass spectrometry (LC-MS/MS). The permeability of the compounds was calculated according to a previous report^57^. The recovery of tested compounds was more than 90%. The permeabilities of Antipyrine (100 μM), Metoprolol (500 μM) and Sulfasarazine (500 μM) as reference compounds, with 100%, 95%, and 13% gastrointestinal absorptions in humans^57^, were 11, 1.5 and 0.055 ’ 10^-6^ cm/s, respectively.

Hepatic microsomal stability assay: Disappearance of the parent compound over time was measured by using the amount of drug at time zero as a reference. After 5 min of preincubation, 1 mM NADPH (final concentration, the same applies to the following) was added to a mixture containing 1 μM of the test compound, 0.2 mg/mL of human or mouse liver microsomes (Sekisui XenoTech LLC), 1 mM EDTA and 0.1 M phosphate buffer (pH 7.4) and incubated at 37 °C for 30 min by rotation at 60 rpm. An aliquot of 50 μL of the incubation mixture was sampled and added to 250 μL of chilled acetonitrile/internal standard (IS). After centrifuging for 15 min at 3,150 × *g* (4°C), the supernatants were diluted with water and analyzed by LC-MS/MS. Hepatic microsomal stability (mL/min/kg, CL_int_) was calculated according to the previous report^58^, using 48.8 (human) or 45.4 (mouse) mg MS protein/g liver and 25.7 (human) or 87.5 (mouse) g liver/kg body weight as scaling factors.

Determination of the unbound fraction in human or mouse plasma: An equilibrium dialysis apparatus was used to determine the unbound fraction for each compound in human or mouse plasma. High Throughput Dialysis Model HTD96b and Dialysis Membrane Strips MWCO 12-14 kDa obtained from HTDialysis, LLC (Gales Ferry, CT) were used. Plasma was spiked with the test compound (1 μM), and 150 μL aliquots were loaded into the apparatus and dialyzed versus 150 μL of 0.1 M phosphate buffer (pH 7.4) at 37°C for 6 h by rotation at 80 rpm. The unbound fraction was calculated as the ratio of receiver side (buffer) to donor side (plasma) concentrations.

### *In vivo* pharmacokinetics assay

Five-week-old female BALB/c mice (purchased from Japan SLC) were treated with s2U by oral (150 mg/kg) or intravenous (20 mg/kg) administration. Blood was collected from the mouse tail with heparin 5 min, 15 min, 30 min, 1 h, 2 h, 4 h, 8 h and 24 h after administration, and plasma samples were isolated at 2,000 rpm for 5 min.

Plasma samples were precipitated with 4–8 volumes of acetonitrile/IS and centrifuged at 15,000 ’ *g* at 4°C for 10 min. The supernatants were diluted with 7 volumes of water and analyzed by LC-MS/MS. Standard non-compartmental analysis was performed to determine the pharmacokinetic parameters and to simulate the repeated dose concentration time profiles using Phoenix Winnonlin ver 8.3 (Pharsight): the estimated initial concentration (C_0_), maximum plasma concentration (C_max_), time to maximum plasma concentration (T_max_), elimination half-life (t_1/2_), area under the concentration time curve from time zero to infinity (AUC_∞_), total clearance (CL_tot_), and volume of distribution at terminal phase (V_dz_). The absolute bioavailability (BA) of the oral dose was calculated as AUC_∞_(po)/AUC_∞_(iv).

### LC-MS/MS quantification method

A Qtrap 6500+ mass spectrometer (Sciex) equipped with a Shimadzu Nexera series LC system (Shimadzu) was used. All compounds were analyzed in multi-reaction monitoring mode under electron spray ionization conditions. The analytical column used was an Acquity UPLC HSS T3 (1.8 μm, 3 ’ 50 mm, Waters) at 40°C. The gradient mobile phase consisted of 0.1% formic acid in water (mobile phase A) and 0.1% formic acid in methanol (mobile phase B) at a total flow rate of 0.5 mL/min. The initial mobile phase composition was 2% B, which was held constant for 0.1 min, increased in a linear fashion to 90% B over 0.9 min, then held constant for 1.5 min, and finally brought back to the initial condition of 2% B over 0.01 min and re-equilibrated for 2.5 min. The transitions (precursor ion > product ion) of s2U and IS (antipyrine) are 261.1 > 129.0 and 189.0 > 56.0 (positive), respectively.

### Statistical analysis

One-way ANOVA followed by Dunnett’s multiple comparisons test or unpaired t test for *in vitro* and *in vivo* antiviral assay and Log–rank (Mantel–Cox) test for *in vivo* survival test were performed to determine the statistical significance using GraphPad Prism version 8.4.3.

## Acknowledgments

We thank Dr. Tomohiko Takasaki (Kanagawa Prefectural Institute of Public Health, Japan) for providing DENV1-4 and CHIKV, Dr. Koichi Morita (Nagasaki University, Japan) for providing SARS-CoV Hanoi strain, Dr. Bart Haagmans (Erasmus University Medical Center, Netherlands) for providing MERS-CoV EMC2012 strain, Drs. Masayuki Saijyo, Masayuki Shimojima, Mutsuyo Ito and Ken Maeda (National Institute of Infectious Diseases (NIID), Japan) for providing SARS-CoV-2 variants, Dr Chang-Kweng Lim (NIID, Japan) for providing RABV HEP strain, Dr. Masayuki Saijo (NIID, Japan) for providing LCMV Armstrong strain, Dr. Akihiro Ishii (Hokkaido University, Japan) for providing LPHV 11SB17 strain, Dr. Shigeru Morikawa (NIID) for providing Junin virus Candid #1 strain, Dr. Shinji Makino (University of Texas Medical Branch, USA) for providing RVFV plasmid, Dr. Karl-Klaus Conzelmann (Max von Pettenkofer-Institute, Germany) for providing BSR-T7/5 cells, Dr. Kumiko Yoshimatsu (Hokkaido University, Japan) for providing TPMV VRC-66412 strain, Dr. Kennedy F. Shortridge (University of Hong Kong, Hong Kong) for providing IAV-H5N1 A/Hong Kong/483/97 strain and Dr. Hiroyuki Miyoshi (RIKEN BRC, Japan) for providing lentiviral vector plasmid CSII-CMV-MCS-IRES2-Bsd. We also thank Takao Sanaki, Yuki Maruyama, Masaaki Izawa, Takao Shishido, Akira Naito, Haruka Maeda, Etsuko Hayashi, Mai Kishimoto, Yukari Itakura, Satoko Otsuguro and Tatsuya Zenko for their excellent assistance. We would like to thank Enago (www.enago.jp) for the English language review. This work is supported by the Scientific Research on Innovative Areas and International Group from the MEXT/JSPS KAKENHI [JP20H05873 (K.Maenaka)], by the Japan Agency for Medical Research and Development (AMED) [JP21wm0225018 (Y.O.), JP22wm0225017 (H.S.), JP223fa627005 (A.S., H.S., K.Maenaka), JP20ae0101047, JP21fk0108463, JP21am0101093, JP22ama121037 (K.Maenaka)], by the Japan Science and Technology Agency (JST) Moonshot R&D [JPMJMS2025 (Y.O., Y.M.)], by the Takeda Foundation (K.Maenaka), and partially supported by the fund from the Atlantic Philanthropies director (W.W.H.).

## Author Contributions

K.U., H.N., A.S., S.T., M.S., Y.O. and H.S. planned and coordinated the experimental virology work. K.U. and M.S. generated the protein expression cells. A.M. and K.Maenaka. synthesized and analyzed the compounds. K.U., H.N., A.S., M.S., K.T. and K.Matsuno. performed *in vitro* antiviral assays. K.U. and H.N. selected drug-resistant mutant and constructed the infectious clones. K.U., S.I., M. T., Y.A., S.Kita. and A.M. performed and analyzed viral RdRp assay. K.U., H.N., S.T. and S.Kusakabe. performed and analyzed *in vivo* experiments. N.M., M.I. and K.K. performed and analyzed a pharmacokinetics study. K.U. wrote the manuscript. A.S., K.K., W.W.H, H.S., A.M. and K.Maenaka. edited the manuscript. A.S., W.W.H., H.S., H.K., A.M. and K.Maenaka. oversaw the conception and supervised the study. Y.O., Y.M., W.W.H., H.S. and K.Maenaka provided the grant support. All authors read and approved the manuscript.

---

**Extended Data Fig. 1 | Identification and cytotoxicity of s2U.**

**a**, Schematic representation of the compound screening using BHK-21 cells and flaviviruses. **b**, Effect of s2U on cell proliferation. Cells were incubated with serial dilutions of the compound. A resazurin reduction assay was performed at 4 days post treatment. Cytotoxicity (%) is expressed relative to the values for the DMSO-treated samples and cell-free samples. The 50% cytotoxic concentration (CC_50_) value was defined in GraphPad Prism version 8.4.3 with a variable slope (four parameters).

**Extended Data Fig. 2 | Dose-response inhibition of several RNA viruses by s2U.**

Cells were infected with DENV2 (multiplicity of infection [MOI] = 0.05), ZIKV (MOI = 0.05), YFV (MOI = 0.05), JEV (MOI = 0.05), WNV (MOI = 0.05), CHIKV (MOI = 0.01), HCoV-229E (MOI = 0.005), HCoV-OC43 (MOI = 0.1), SARS-CoV (MOI = 0.01), MERS-CoV (MOI = 0.01) and several SARS-CoV-2 variants (MOI = 0.01) containing a serially diluted compound. Cell lysates were collected for viral RNA determination, and viral RNA levels were determined relative to *ACTB* or *18S rRNA* transcripts. The 90% effective concentration (EC_90_) value was defined in GraphPad Prism version 8.4.3 with a variable slope (Find ECanything; F = 90). Data are presented as mean values of biological triplicates from one of the experiments, and error bars indicate SD.

**Extended Data Fig. 3 | Molecular target and mechanism of action of s2U (related to Fig. 2).**

**a**, Ribonucleotide competition for HCoV-229E inhibition by s2U. HCoV-229E (MOI = 0.005)-infected MRC5 cells were treated with 15 μM of s2U and serial dilutions of exogenous nucleosides. A resazurin reduction assay was performed at 3 days post infection (dpi). Antiviral activities (%) are expressed relative to the values for the DMSO-treated, infected samples and non-infected samples. **b, c**, Effect of s2U resistance mutation on replication fitness. BHK-21 (**b**) and VeroE6 (**c**) cells were infected with rgDENV2-WT or rgDENV2-NS5-G605V (MOI = 0.01) for 1 h. Cell lysates were collected at 24, 48, 72 and 96 hpi, and viral RNA levels were determined relative to *18S rRNA* (BHK-21) or *ACTB* (VeroE6) transcripts. **d**, Effect of s2U resistance mutation on anti-DENV2 activity of s2U. VeroE6 cells were infected with rgDENV2-WT or rgDENV2-NS5-G605V (MOI = 0.05) containing a serially diluted compound. Cell lysates were collected at 72 hpi, and viral RNA levels were determined relative to *ACTB* transcripts. Inhibitory effect (% Inhibition) is expressed relative to the values for the DMSO-treated samples. Data are presented as mean values, and error bars indicate SD.

**Extended Data Fig. 4 | Effect of s2U on mitochondrial biogenesis.**

**a–d**, HepG2 cells were assayed for a reduction in mitochondrial-encoded protein COX-I or nuclear-encoded protein SDH-A after 5 days of incubation with 3-fold serial dilutions of s2U (**a**), ribavirin (**b**), favipiravir (**c**) and chloramphenicol (**d**). Inhibitory effects (% of Control) are expressed relative to the values for the DMSO-treated samples. **d**, 50% inhibitory concentration (IC_50_) values of these compounds against protein expression. The IC_50_ value was defined in GraphPad Prism version 8.4.3 with a variable slope (four parameters).

**Extended Data Fig. 5 | Establishment of mouse-adapted DENV2 strain (DENV2 AG-P10).**

**a**, Schematic representation of the passage history of DENV2 in AG129 mice. Virus in serum from infected mice was propagated in C6/36 cells. **b**, Survival of DENV2 AG-P10-infected AG129 mice. Mice were intraperitoneally inoculated with 4 × 10^5^ plaque forming units [PFU] of a DENV2 clinical isolate (n = 3) and DENV2 AG-P10 (n = 5). Survival was monitored daily. **c**, Viral RNA copies/mL in organ samples were quantified using qRT-PCR. At 4 dpi, the infected mice (1 × 10^3^ PFU of DENV2 AG-P10, n = 2) were euthanized under deep anesthesia by isoflurane inhalation, and serum and whole tissues (spleen, kidney, liver, small intestine, large intestine and brain) were harvested and homogenized in PBS with a TissueRuptor. **d**, Amino acid substitutions occurred during passage. Data are presented as mean values, and error bars indicate SD.

**Extended Data Fig. 6 | Establishment of mouse-adapted SARS-CoV-2 strain (SARS-CoV-2 MA-P10).**

**a**, Schematic representation of the passage history of SARS-CoV-2 in BALB/c mice. **b**, Virus titers in lung homogenates from SARS-CoV-2-infected mice from passage 1 (P1) to P10 (n = 3–9). **c**, Survival of SARS-CoV-2 MA-P10-infected BALB/c mice. Young (5-week-old) and adult (30–50-week-old) female mice were intranasally inoculated with 2 × 10^5^ TCID_50_ of SARS-CoV-2 MA-P10 (n = 5 per group). Survival was monitored daily. **d, e**, Virus titers and viral RNA loads in lung from SARS-CoV-2-MA-P10-infected mice. Virus titers (**d**) were quantified by standard 50% tissue culture infection dose (TCID_50_) assay using VeroE6/TMPRSS2 cells. Viral RNA copies/mL (**e**) were quantified using qRT-PCR. **f**, Macroscopic appearance of lung tissue of SARS-CoV-2-MA-P10-infected mice at 1, 3 and 5 dpi. **g**, Amino acid substitutions occurred during passage. Data are presented as mean values, and error bars indicate SD.

**Extended Data Fig. 7 | *In vivo* efficacy of s2U in the SARS-CoV-2 mouse model (related to Fig. 3).**

**a**, Schematic representation of the survival and viremia studies using BALB/c mice and SARS-CoV-2 MA-P10. **b**, Effect of s2U on viremia at 1 dpi in mice orally administered 300 mg/kg s2U twice daily compared with vehicle-treated mice (n = 5 per group). Virus titers in lung samples were quantified by a standard TCID_50_ assay using VeroE6/TMPRSS2 cells. Data are presented as mean values, and error bars indicate SD. Statistically significant differences were determined using the unpaired *t*-test to compare s2U-treated with vehicle-treated mice; \*\*\*\**p* < 0.0001.

**Extended Data Fig. 8 | Pharmacokinetic (PK) properties of s2U.**

**a**, *In vitro* absorption, distribution, metabolism and excretion (ADME) properties of s2U. **b–d**, Pharmacokinetic properties of s2U in mice after oral (**b**) and intravenous (**c**) dosing. s2U was administered to 5-week-old female BALB/c mice (n = 3 per group) *via* oral gavage as a solution formulated in 5% DMSO/0.5% methylcellulose at 150 mg/kg or intravenously as a saline solution at 20 mg/kg. C_0_: initial concentration, C_max_: maximum plasma concentration, Tmax: time to reach C_max_, t_1/2_: terminal phase elimination half-life, AUC: area under the plasma concentration versus the time, AUC_∞_: AUC curve to infinite time, CL_tot_: total clearance, V_dz_: volume of distribution at the terminal phase, BA (F): bioavailability. **e, f**, Simulation of twice-daily or once-daily doses of s2U by oral or intravenous administration derived from the single-dose PK experiment. Data are presented as mean values, and error bars indicate SD (**b, c**).

**Extended Data Table 1 | Antiviral activity of reference compounds against various RNA viruses.**

Antiviral assays were carried out as described in Supplementary Table 1. EC_50_: 50% effective concentration. EC_90_: 90% effective concentration.

a: EC_50_ values represent mean values from at least three independently performed experiments (n = 2).

b: EC_90_ values represent mean values from a single experiment with biological triplicates. c: Fold change is calculated from the ratio of rgNS5-G605V/rgWT.

**Supplementary Table 1 | *In vitro* assay conditions.**

**Supplementary Table 2 | Sequence of primers and probes for qPCR assay.**

