## Supplementary materials for "2-Thiouridine is a broad-spectrum antiviral nucleoside analogue against positive-strand RNA viruses"

### Extended Data Figure 1

**a**

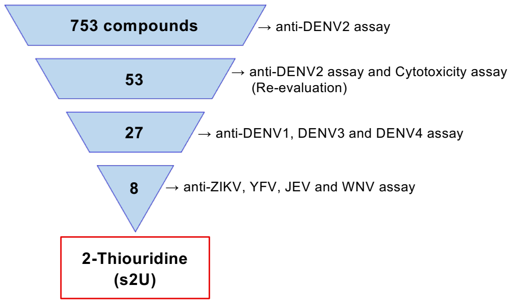


**b**

| Cell | Species | Organ | $CC_{50}$ (μM) [a] |
|---|---|---|---|
| BHK-21 | Hamster | Kidney | > 400 [b,c] |
| VeroE6 | Monkey | Kidney | > 400 [b,c] |
| MRC5 | Human | Lung | > 400 [b] |
| 293T | Human | Kidney | 100.6 [c] |
| MOLT4 | Human | T cell | > 50 [c] |
| THP-1 | Human | Monocyte | > 50 [c] |

a : $CC_{50}$ values represent mean values from at least three independently performed experiments (n = 2).
b : CellTiter-Glo assay
c : Resazurin assay

**Extended Data Fig. 1 | Identification and cytotoxicity of s2U.**
**a**, Schematic of the compound screening using BHK-21 cells and flaviviruses. **b**, Effect of s2U on cell proliferation. Cells were incubated with serial dilutions of the compound. A resazurin reduction assay was performed at 4 days post treatment. Cytotoxicity (%) is expressed relative to the values for the DMSO-treated samples and cell-free samples. The 50% cytotoxic concentration ($CC_{50}$) value was defined in GraphPad Prism version 8.4.3 with a variable slope (four parameters).

**Extended Data Figure 2**

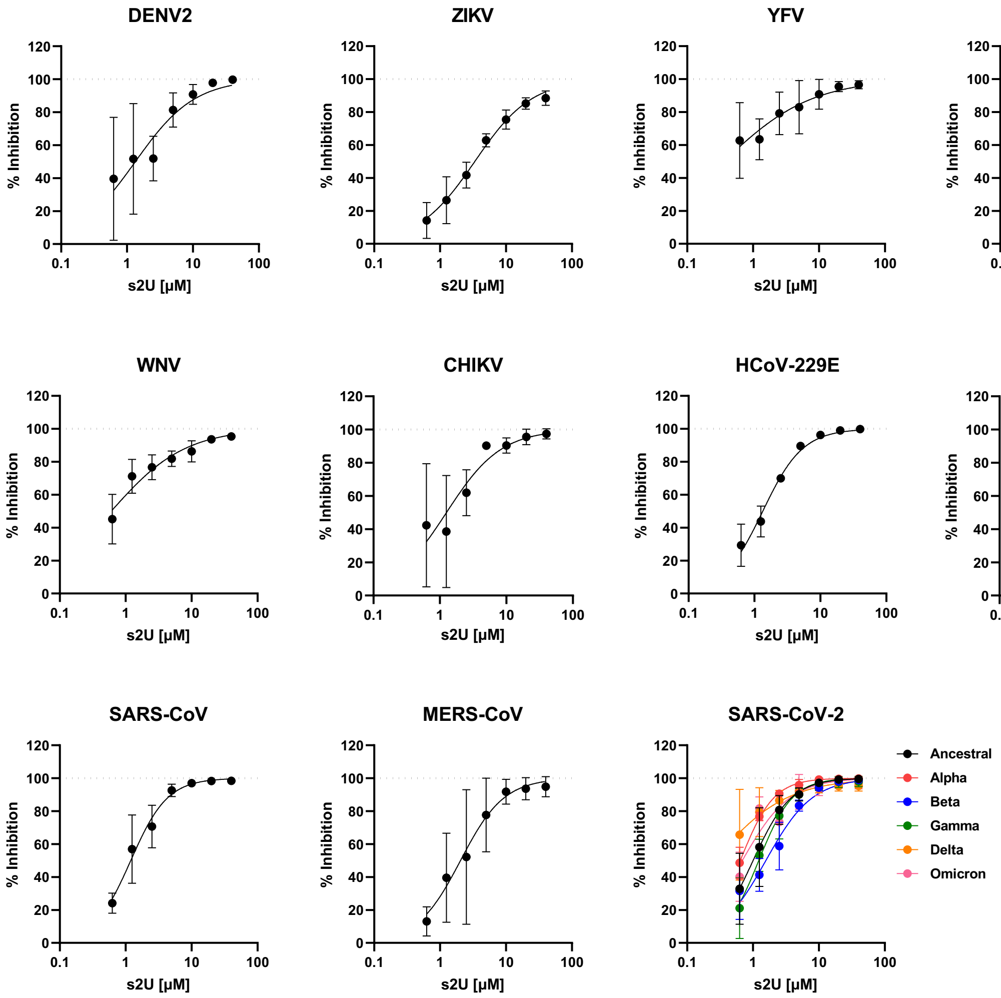


**Extended Data Fig. 2 | Dose-response inhibition of several RNA viruses by s2U.**
Cells were infected with DENV2 (multiplicity of infection [MOI] = 0.05), ZIKV (MOI = 0.05), YFV (MOI = 0.05), JEV (MOI = 0.05), WNV (MOI = 0.05), CHIKV (MOI = 0.01), HCoV-229E (MOI = 0.005), HCoV-OC43 (MOI = 0.1), SARS-CoV (MOI = 0.01), MERS-CoV (MOI = 0.01) and several SARS-CoV-2 variants (MOI = 0.01) containing a serially diluted compound. Cell lysates were collected for viral RNA determination, and viral RNA levels were determined relative to *ACTB* or *18S rRNA* transcripts. The 90% effective concentration ($EC_{90}$) value was defined in GraphPad Prism version 8.4.3 with a variable slope (Find ECanything; F = 90). Data are presented as mean values of biological triplicates from one of the experiments, and error bars indicate SD.

#### Extended Data Figure 3

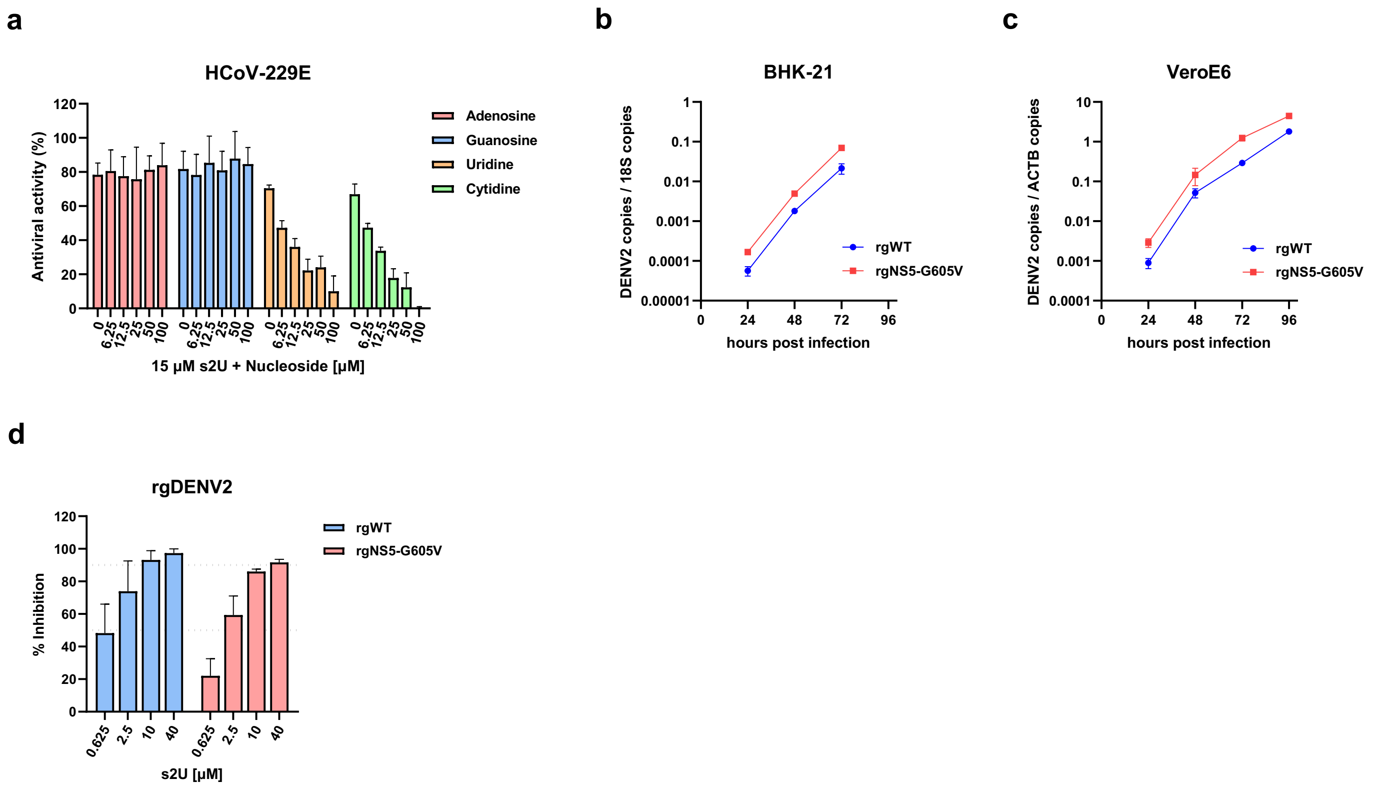


**Extended Data Fig. 3 | Molecular target and mechanism of action of s2U (related to Fig. 2).** **a**, Ribonucleotide competition for HCoV-229E inhibition by s2U. HCoV-229E (MOI = 0.005)-infected MRC5 cells were treated with 15 µM of s2U and serial dilutions of exogenous nucleosides. A resazurin reduction assay was performed at 3 days post infection (dpi). Antiviral activities (%) are expressed relative to the values for the DMSO-treated, infected samples and non-infected samples. **b, c**, Effect of s2U resistance mutation on replication fitness. BHK-21 (**b**) and VeroE6 (**c**) cells were infected with rgDENV2-WT or rgDENV2-NS5-G605V (MOI = 0.01) for 1 h. Cell lysates were collected at 24, 48, 72 and 96 hpi, and viral RNA levels were determined relative to *18S rRNA* (BHK-21) or *ACTB* (VeroE6) transcripts. **d**, Effect of s2U resistance mutation on anti-DENV2 activity of s2U. VeroE6 cells were infected with rgDENV2-WT or rgDENV2-NS5-G605V (MOI = 0.05) containing a serially diluted compound. Cell lysates were collected at 72 hpi, and viral RNA levels were determined relative to *ACTB* transcripts. Inhibitory effect (% Inhibition) is expressed relative to the values for the DMSO-treated samples. Data are presented as mean values, and error bars indicate SD.

### Extended Data Figure 4

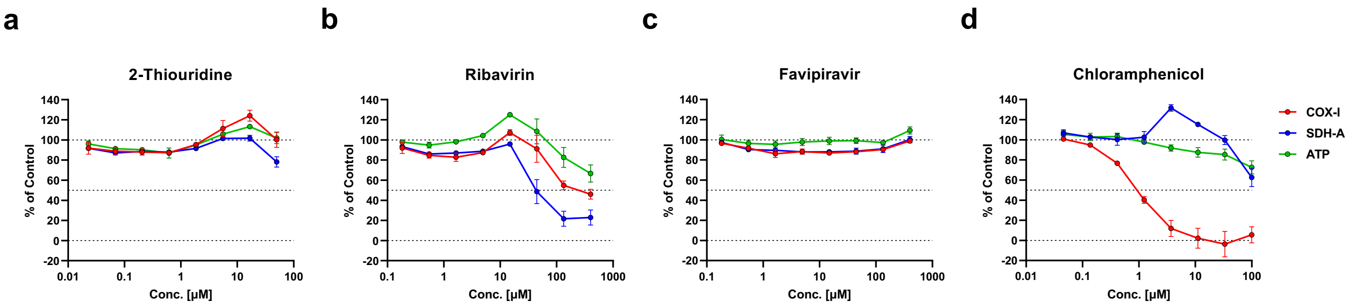


e

| Protein | IC50 (μM) [a] 2-Thiouridine | Ribavirin | Favipiravir | Chloramphenicol [e] |
|---|---|---|---|---|
| COX-I [b] | >50 | 97 ± 33 | >400 | 0.97 ± 0.060 |
| SDH-A [c] | >50 | 45 ± 1.4 | >400 | >100 |
| ATP [d] | >50 | >400 | >400 | >100 |

a : $IC_{50}$ values represent mean values ± SEM from three independently performed experiments (n = 2).
b : Cytochrome *c* oxidase (COX-I) is the mitochondrial DNA-encoded protein.
c : Succinate dehydrogenase A subunit (SDH-A) is the nuclear DNA-encoded protein.
d : Cell viability monitored as ATP level was measured using Cell-Titer Glo® 2.0 Assay.
e : Chloramphenicol was used as control compound for specific inhibition of COX-I synthesis.

**Extended Data Fig. 4 | Effect of s2U on mitochondrial biogenesis.**
**a–d**, HepG2 cells were assayed for a reduction in mitochondrial-encoded protein COX-I or nuclear-encoded protein SDH-A after 5 days of incubation with 3-fold serial dilutions of s2U (**a**), ribavirin (**b**), favipiravir (**c**) and chloramphenicol (**d**). Inhibitory effects (% of Control) are expressed relative to the values for the DMSO-treated samples. **e**, 50% inhibitory concentration ($IC_{50}$) values of these compounds against protein expression. The $IC_{50}$ value was defined in GraphPad Prism version 8.4.3 with a variable slope (four parameters).

#### Extended Data Figure 5

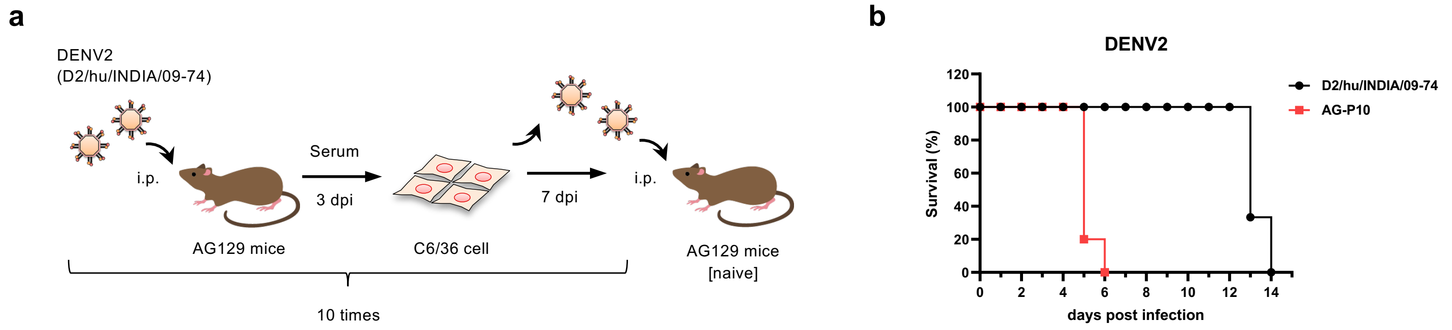


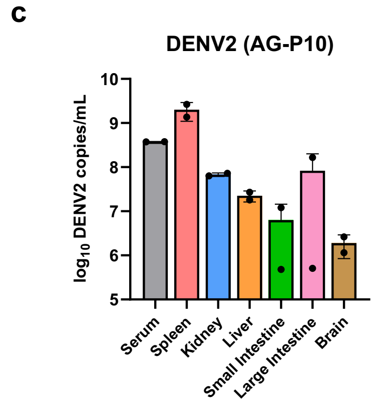


**d**

| Viral protein | NS4B | NS5 |
|---|---|---|
| Amino acid substitution | A119T | E802Q |

**Extended Data Fig. 5 | Establishment of mouse-adapted DENV2 strain (DENV2 AG-P10).**
**a**, Schematic of the passage history of DENV2 in AG129 mice. Virus in serum from infected mice was propagated in C6/36 cells. **b**, Survival of DENV2 AG-P10-infected AG129 mice. Mice were intraperitoneally inoculated with $4 \times 10^5$ plaque forming units [PFU] of a DENV2 clinical isolate (n = 3) and DENV2 AG-P10 (n = 5). Survival was monitored daily. **c**, Viral RNA copies/mL in organ samples were quantified using qRT-PCR. At 4 dpi, the infected mice ($1 \times 10^3$ PFU of DENV2 AG-P10, n = 2) were euthanized under deep anesthesia by isoflurane inhalation, and serum and whole tissues (spleen, kidney, liver, small intestine, large intestine and brain) were harvested and homogenized in PBS with a TissueRuptor. **d**, Amino acid substitutions occurred during passage. Data are presented as mean values, and error bars indicate SD.

#### Extended Data Figure 6

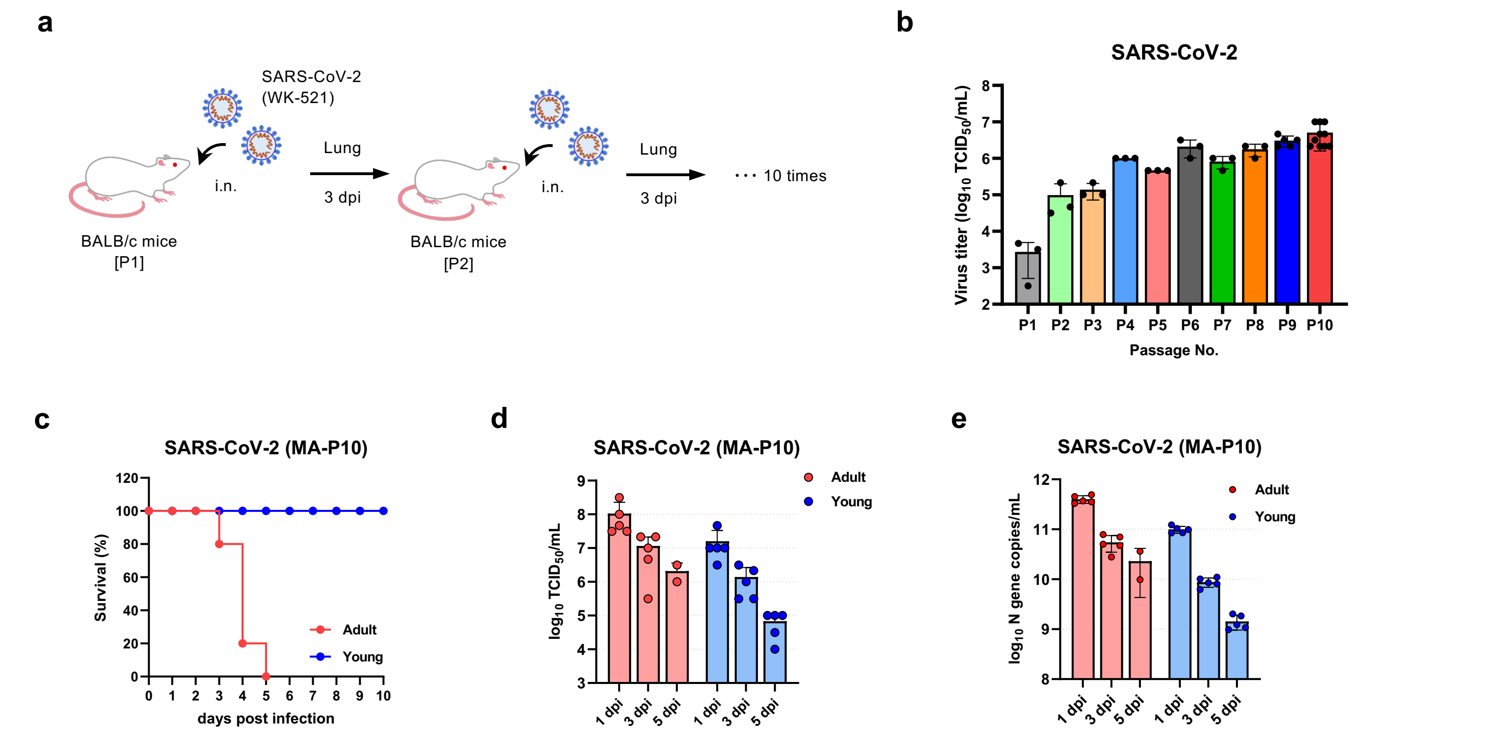


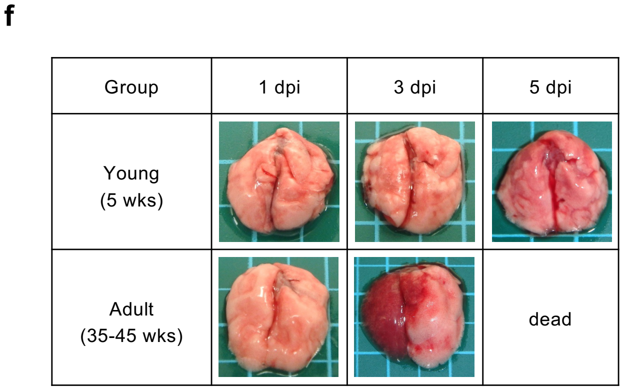


**g**

| Viral protein | Amino acid substitution |
|---|---|
| nsp4 | A457V |
| nsp5 | E270G |
| nsp9 | R39K |
| nsp16 | K160R |
| Spike | Q498H |
| Nucleocapsid | T205I |

**Extended Data Fig. 6 | Establishment of mouse-adapted SARS-CoV-2 strain (SARS-CoV-2 MA-P10).** **a**, Schematic of the passage history of SARS-CoV-2 in BALB/c mice. **b**, Virus titers in lung homogenates from SARS-CoV-2-infected mice from passage 1 (P1) to P10 (n = 3–9). **c**, Survival of SARS-CoV-2 MA-P10-infected BALB/c mice. Young (5-week-old) and adult (30–50-week-old) female mice were intranasally inoculated with $2 \times 10^5$ $TCID_{50}$ of SARS-CoV-2 MA-P10 (n = 5 per group). Survival was monitored daily. **d, e**, Virus titers and viral RNA loads in lung from SARS-CoV-2-MA-P10-infected mice. Virus titers (**d**) were quantified by standard 50% tissue culture infection dose ($TCID_{50}$) assay using VeroE6/TMPRSS2 cells. Viral RNA copies/mL (**e**) were quantified using qRT-PCR. **f**, Macroscopic appearance of lung tissue of SARS-CoV-2-MA-P10-infected mice at 1, 3 and 5 dpi. **g**, Amino acid substitutions occurred during passage. Data are presented as mean values, and error bars indicate SD.

### Extended Data Figure 7

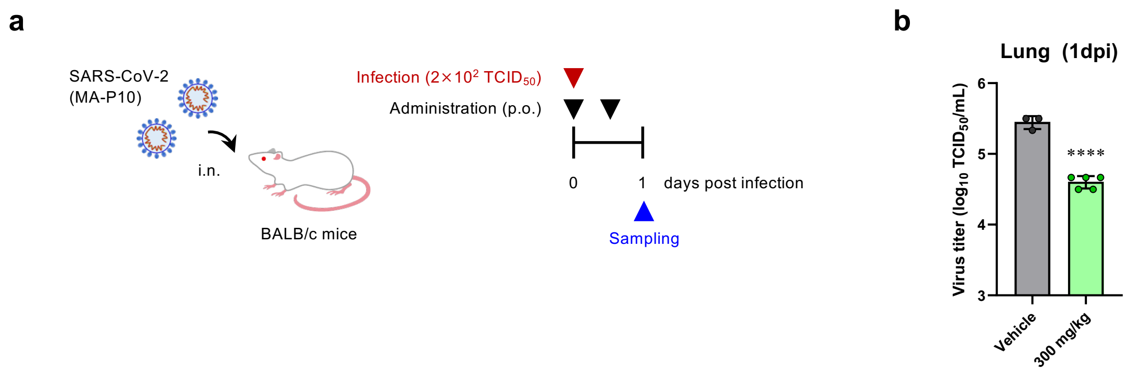


**Extended Data Fig. 7 | *In vivo* efficacy of s2U in the SARS-CoV-2 mouse model (related to Fig. 3).** **a**, Schematic of the survival and viremia studies using BALB/c mice and SARS-CoV-2 MA-P10. **b**, Effect of s2U on viremia at 1 dpi in mice orally administered 300 mg/kg s2U twice daily compared with vehicle-treated mice (n = 5 per group). Virus titers in lung samples were quantified by a standard TCID$_{50}$ assay using VeroE6/TMPRSS2 cells. Data are presented as mean values, and error bars indicate SD. Statistically significant differences were determined using the unpaired *t*-test to compare s2U-treated with vehicle-treated mice; **** $p < 0.0001$.

### Extended Data Figure 8

**a**

| Compound | Solubility (μM) | | Membrane Permeability ($\times 10^{-6}$ cm/sec) | Metabolic stability (mL/min/kg) | | Protein binding (Unbound fraction) | |
|---|---|---|---|---|---|---|---|
| | JP1 (pH1.2) | JP2 (pH6.8) | PAMPA (pH6.5) | Human liver microsome | Mouse liver microsome | Human plasma | Mouse plasma |
| 2-Thiouridine | 78 | >100 | 0.013 | <22.0 (<17.6μL/min/mg) | <69.8 (<17.6μL/min/mg) | 0.70 | 0.73 |

**b**

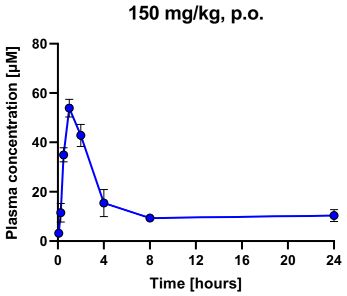


**c**

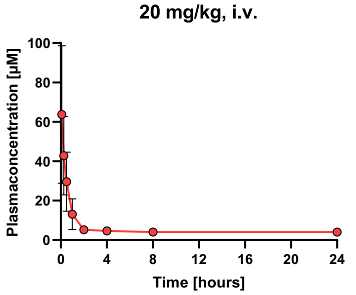


**d**

| Compound | Route | Dose (mg/kg) | $C_0$ (μg/mL) | $C_{max}$ (μg/mL) | $T_{max}$ (hr) | $t_{1/2}$ (hr) | $AUC_\infty$ (μg/mL·hr) | $CL_{tot}$ or $CL_{tot}/F$ (mL/hr/kg) | $V_{dz}$ or $V_{dz}/F$ (mL/kg) | BA (F) (%) |
|---|---|---|---|---|---|---|---|---|---|---|
| 2-Thiouridine | p.o. | 150 | | 14.0 | 1 | 31.2 | 198 | 757 | 34025 | 17.3 |
| | i.v. | 20 | 20 | | | 80.7 | 153 | 131.1 | 15260 | |

**e**

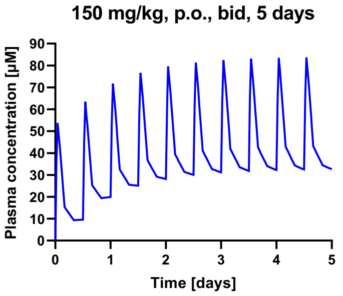


**f**

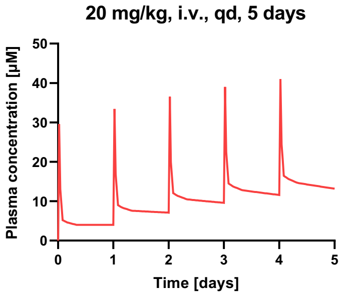


**Extended Data Fig. 8 | Pharmacokinetic (PK) properties of s2U.**
**a**, *In vitro* absorption, distribution, metabolism and excretion (ADME) properties of s2U. **b–d**, Pharmacokinetic properties of s2U in mice after oral (**b**) and intravenous (**c**) dosing. s2U was administered to 5-week-old female BALB/c mice (n = 3 per group) *via* oral gavage as a solution formulated in 5% DMSO/0.5% methyl cellulose at 150 mg/kg or intravenously as a saline solution at 20 mg/kg. $C_0$: initial concentration, $C_{max}$: maximum plasma concentration, $T_{max}$: time to reach $C_{max}$, $t_{1/2}$: terminal phase elimination half-life, AUC: area under the plasma concentration versus the time, $AUC_\infty$: AUC curve to infinite time, $CL_{tot}$: total clearance, $V_{dz}$: volume of distribution at the terminal phase, BA (F): bioavailability. **e, f**, Simulation of twice-daily or once-daily doses of s2U by oral or intravenous administration derived from the single-dose PK experiment. Data are presented as mean values, and error bars indicate SD (**b, c**).

### Extended Data Table 1

#### Extended Data Table 1 | Antiviral activity of reference compounds against various RNA viruses.

| MTT / Resazurin Assay Results | | | $EC_{50}$ (μM) [a] | | | |
|---|---|---|---|---|---|---|
| Virus | Strain | Cell | Ribavirin | Favipiravir | Remdesivir | GS-441524 |
| DENV1 | D1/hu/PHL/10-07 | BHK-21 | 46 | 18 | | |
| DENV2 | D2/hu/INDIA/09-74 | BHK-21 | 32 | 20 | | |
| DENV3 | D3/hu/Thailand/00-40 | BHK-21 | 61 | 13 | | |
| DENV4 | D4/hu/Solomon/09–11 | BHK-21 | 91 | 5.2 | | |
| ZIKV | MR766 | BHK-21 | 18 | 39 | | |
| YFV | 17D-204 | BHK-21 | 15 | 24 | | |
| JEV | Beijing-1 | BHK-21 | 33 | 80 | | |
| WNV | NY99 | BHK-21 | 43 | 36 | | |
| CHIKV | SL10571 | BHK-21 | 46 | 8.1 | | |
| HCoV | 229E | MRC5 | | | 0.071 | 0.76 |
| HCoV | OC43 | MRC5 | | | 0.23 | 2.0 |
| RABV | HEP | BHK-21 | 16 | 5.9 | | |
| LACV | ATCC VR-1834 | MDBK | 2.5 | 20 | | |
| LPHV | 11SB17 | KB | 5.3 | >100 | | |
| LCMV | Armstrong | KB | 1.8 | 21 | | |
| JUNV | Candid #1 | 293T | 2.6 | 10 | | |
| SFTSV | ArtLN/2017 | MDCK | 5.4 | 7.3 | | |
| RVFV | MP12 | MDCK | 17 | 21 | | |
| TPMV | VRC-66412 | VeroE6 | 6.7 | 31 | | |
| IAV H5N1 | A/Hong Kong/483/97 | A549 | 26 | 6.2 | | |
| IAV H7N9 | A/Anhui/1/2013 | MA104/TMPRSS2 | 3.7 | 12 | | |
| **Resazurin Assay Results** | | | $EC_{50}$ (μM) [a] | | Fold change [c] | |
| Virus | Strain | Cell | Ribavirin | Favipiravir | Ribavirin | Favipiravir |
| rgDENV2 | Wild type | BHK-21 | 16 | 7.9 | | |
| rgDENV2 | NS5-G605V | BHK-21 | 37 | 17 | 2.3 | 2.2 |
| **qPCR Assay Results** | | | $EC_{90}$ (μM) [b] | | | |
| Virus | Strain | Cell | Remdesivir | | | |
| SARS-CoV | Hanoi | VeroE6 | 2.7 | | | |
| SARS-CoV-2 | WK-521 (Ancestral) | VeroE6/TMPRSS2 | 3.3 | | | |

a : $EC_{50}$ values represent mean values from at least three independently performed experiments (n = 2).

b : $EC_{90}$ values represent mean values from a single experiment with biological triplicates.

c : Fold change is calculated from the ratio of rgNS5-G605V/rgWT.

### Supplementary Table 1

#### Supplementary Table 1 | *In vitro* assay conditions.

**MTT / Resazurin Assay**

| Virus | Strain | Cell | MOI | Method | Period |
|---|---|---|---|---|---|
| DENV1 | D1/hu/PHL/10-07 | BHK-21 | 10 $TCID_{50}$ | Resazurin | 5 days |
| DENV2 | D2/hu/INDIA/09-74 | BHK-21 | MOI=0.01 | Resazurin | 4 days |
| DENV3 | D3/hu/Thailand/00-40 | BHK-21 | 10 $TCID_{50}$ | Resazurin | 5 days |
| DENV4 | D4/hu/Solomon/09–11 | BHK-21 | 10 $TCID_{50}$ | Resazurin | 5 days |
| ZIKV | MR766 | BHK-21 | MOI=0.01 | MTT | 3 days |
| YFV | 17D-204 | BHK-21 | MOI=0.01 | MTT | 3 days |
| JEV | Beijing-1 | BHK-21 | 10 $TCID_{50}$ | MTT | 3 days |
| | | VeroE6 | MOI=0.01 | MTT | |
| WNV | NY99 | BHK-21 | 10 $TCID_{50}$ | MTT | 3 days |
| | | VeroE6 | MOI=0.01 | MTT | |
| CHIKV | SL10571 | BHK-21 | 10 $TCID_{50}$ | MTT | 3 days |
| | | VeroE6 | MOI=0.01 | MTT | |
| HCoV | 229E | MRC5 | MOI=0.005 | Resazurin | 3 days |
| HCoV | OC43 | MRC5 | MOI=0.01 | Resazurin | 3 days |
| RABV | HEP | BHK-21 | 10 $TCID_{50}$ | MTT | 3 days |
| LACV | ATCC VR-1834 | MDBK | 10 $TCID_{50}$ | MTT | 3 days |
| LPHV | 11SB17 | KB | 10 $TCID_{50}$ | MTT | 4 days |
| LCMV | Armstrong | KB | 10 $TCID_{50}$ | MTT | 4 days |
| JUNV | Candid #1 | 293T | 10 $TCID_{50}$ | MTT | 4 days |
| SFTSV | ArtLN/2017 | MDCK | 10 $TCID_{50}$ | MTT | 4 days |
| RVFV | MP12 | MDCK | 10 $TCID_{50}$ | MTT | 3 days |
| TPMV | VRC-66412 | VeroE6 | 10 $TCID_{50}$ | MTT | 4 days |
| IAV H5N1 | A/Hong Kong/483/97 | A549 | 10 $TCID_{50}$ | MTT | 3 days |
| IAV H7N9 | A/Anhui/1/2013 | MA104/TMPRSS2 | 10 $TCID_{50}$ | MTT | 3 days |
| rgDENV2 | Wild type | BHK-21 | MOI=0.1 | Resazurin | 4 days |
| rgDENV2 | NS5-G605V | BHK-21 | MOI=0.1 | Resazurin | 4 days |

**qRT-PCR (qPCR) Assay**

| Virus | Strain | Cell | MOI | Period | Internal control |
|---|---|---|---|---|---|
| DENV2 | D2/hu/INDIA/09-74 | VeroE6 | MOI=0.05 | 48 hpi | ACTB |
| ZIKV | MR766 | VeroE6 | MOI=0.05 | 48 hpi | ACTB |
| YFV | 17D-204 | VeroE6 | MOI=0.05 | 48 hpi | ACTB |
| JEV | Beijing-1 | VeroE6 | MOI=0.01 | 48 hpi | ACTB |
| WNV | NY99 | VeroE6 | MOI=0.05 | 24 hpi | ACTB |
| CHIKV | SL10571 | VeroE6 | MOI=0.01 | 24 hpi | ACTB |
| HCoV | 229E | MRC5 | MOI=0.005 | 48 hpi | ACTB |
| HCoV | OC43 | VeroE6 | MOI=0.1 | 48 hpi | ACTB |
| SARS-CoV | Hanoi | VeroE6/TMPRSS2 | MOI=0.01 | 24 hpi | ACTB |
| MERS-CoV | EMC2012 | VeroE6/TMPRSS2 | MOI=0.01 | 24 hpi | ACTB |
| SARS-CoV-2 | WK-521 (Ancestral) | VeroE6/TMPRSS2 | MOI=0.01 | 24 hpi | ACTB |
| | QK002 (Alpha) | VeroE6/TMPRSS2 | MOI=0.01 | 24 hpi | ACTB |
| | TY8-612 (Beta) | VeroE6/TMPRSS2 | MOI=0.01 | 24 hpi | ACTB |
| | TY7-501 (Gamma) | VeroE6/TMPRSS2 | MOI=0.01 | 24 hpi | ACTB |
| | TY11-927 (Delta) | VeroE6/TMPRSS2 | MOI=0.01 | 24 hpi | ACTB |
| | TY38-873 (Omicron) | VeroE6/TMPRSS2 | MOI=0.01 | 24 hpi | ACTB |
| RABV | HEP | BHK-21 | MOI=0.1 | 24 hpi | 18S rRNA |
| RVFV | MP12 | VeroE6 | MOI=0.01 | 24 hpi | ACTB |
| HSV-1 | F | VeroE6 | MOI=0.1 | 24 hpi | ACTB |
| rgDENV2 | Wild type | VeroE6 | MOI=0.05 | 72 hpi | ACTB |
| rgDENV2 | NS5-G605V | VeroE6 | MOI=0.05 | 72 hpi | ACTB |

**IFA**

| Virus | Strain | Cell | MOI | Period |
|---|---|---|---|---|
| DENV2 | D2/hu/INDIA/09-74 | VeroE6 | MOI=0.1 | 48 hpi |
| CHIKV | SL10571 | VeroE6 | MOI=0.05 | 24 hpi |
| HCoV | OC43 | VeroE6/TMPRSS2 | MOI=0.1 | 48 hpi |
| SARS-CoV-2 | WK-521 (Ancestral) | VeroE6/TMPRSS2 | MOI=0.005 | 24 hpi |

### Supplementary Table 2

#### Supplementary Table 2 | Sequence of primers and probes for qPCR assay.

| Virus | | Sequence (5'→3') |
|---|---|---|
| DENV2 | Fw | AGTGGACACGAGAACCCAAGA |
| | Rv | TTCGGCCGTGATTTTCATTAG |
| | Pr | FAM/AAAAGAAGGCACGAAGAA/MGB |
| ZIKV | Fw | GACATGGCTTCGGACAG |
| | Rv | CTTTGCCAAAAAGTCCACA |
| YFV | Fw | GCTAATTGAGGTGYATTGGTCTGC |
| | Rv | CTGCTAATCGCTCAAMGAACG |
| | Pr | 56-FAM/ATCGAGTTG/ZEN/CTAGGCAATAAACAC/3IABkFQ |
| JEV | Fw | AGCTGGGCCTTCTGGT |
| | Rv | CCCAAGCATCAGCACAAG |
| | Pr | 56-FAM/CTTCGCAAG/ZEN/AGGTGGACGGCCA/3IABkFQ |
| WNV | Fw | AAGTTGAGTAGACGGTGCTG |
| | Rv | AGACGGTTCTGAGGGCTTAC |
| | Pr | 56-FAM/GCTCAACCCCAGGAGGACTGG/MGBEQ |
| CHIKV | Fw | AAGCTYCGCGTCCTTTACCAAG |
| | Rv | CCAAATTGTCCYGGTCTTCCT |
| | Pr | 56-FAM/CCAATGTCT/ZEN/TCAGCCTGGACACCTTT/3IABkFQ |
| HCoV-229E | Fw | CAGTCAAATGGGCTGATGCA |
| | Rv | CAAAGGGCTATAAAGAGAATAAGGTATTCT |
| | Pr | 56-FAM/TGAACCACA/ZEN/ACGTGGTCGTCAGGG/3IABkFQ |
| HCoV-OC43 | Fw | CGATGAGGCTATTCCGACTAGGT |
| | Rv | CTTCCTGAGCCTTCAATATAGTAACC |
| | Pr | 56-FAM/TCCGCCTGG/ZEN/CACGGTACTCCCT/3IABkFQ |
| SARS-CoV | Fw | GGAGCCTTGAATACACCCAAAG |
| | Rv | GCACGGTGGCAGCATTG |
| | Pr | 56-FAM/CCACATTGG/ZEN/CACCCGCAATCC/3IABkFQ |
| MERS-CoV | Fw | CAAAACCTTCCCTAAGAAGGAAAAG |
| | Rv | GCTCCTTTGGAGGTTCAGACAT |
| | Pr | 56-FAM/ACAAAAGGC/ZEN/ACCAAAAGAAGAATCAACAGACC/3IABkFQ |
| SARS-CoV-2 | Fw | CACATTGGCACCCGCAATC |
| | Rv | GAGGAACGAGAAGAGGCTTG |
| | Pr | 56-FAM/ACTTCCTCA/ZEN/AGGAACAACATTGCCA/3IABkFQ |
| RABV | Fw | GCCACGGTTATTGCTGCAT |
| | Rv | CTCCCAAATAGCCCCCTAGAA |
| | Pr | FAM/CCCTCATGAGATGTC/MGB |
| RVFV | Fw | TTCTTTGCTTCTGATACCCTCTG |
| | Rv | GTTCCACTTCCTTGCATCATCTG |
| | Pr | 56-FAM/TTGCACAAG/ZEN/TCCACACAGGCCCCT/3IABkFQ |
| HSV-1 | Fw | GGGCCGTGATTTTGTTTGTC |
| | Rv | CCGCCAAGGCATATTTGC |
| | Pr | FAM/TAGTGGGCCTCCATGGG/MGB |
| ACTB | Fw | GCTGCCCTGAGGCTCTCTT |
| (*C.aethiops*) | Rv | TGATGGAGTTGAAGGTAGTTTCATG |
